# Sequential emergence and contraction of epithelial subtypes in the prenatal human choroid plexus revealed by a stem cell model

**DOI:** 10.1101/2024.06.12.598747

**Authors:** Haley Masters, Shuxiong Wang, Christina Tu, Quy Nguyen, Yutong Sha, Matthew K. Karikomi, Pamela Shi Ru Fung, Benjamin Tran, Cristina Martel, Nellie Kwang, Michael Neel, Olga G. Jaime, Victoria Espericueta, Brett A. Johnson, Kai Kessenbrock, Qing Nie, Edwin S. Monuki

## Abstract

Despite the major roles of choroid plexus epithelial cells (CPECs) in brain homeostasis and repair, their developmental lineage and diversity remain undefined. In simplified differentiations from human pluripotent stem cells, derived CPECs (dCPECs) displayed canonical properties and dynamic multiciliated phenotypes that interacted with Aβ uptake. Single dCPEC transcriptomes over time correlated well with human organoid and fetal CPECs, while pseudotemporal and cell cycle analyses highlighted the direct CPEC origin from neuroepithelial cells. In addition, time series analyses defined metabolic (type 1) and ciliogenic dCPECs (type 2) at early timepoints, followed by type 1 diversification into anabolic-secretory (type 1a) and catabolic-absorptive subtypes (type 1b) as type 2 cells contracted. These temporal patterns were then confirmed in independent derivations and mapped to prenatal stages using human tissues. In addition to defining the prenatal lineage of human CPECs, these findings suggest new dynamic models of ChP support for the developing human brain.

## INTRODUCTION

Inside the brain’s ventricles, the choroid plexus (ChP) lies at the interface between two circulating fluids^1^ – the blood and cerebrospinal fluid (CSF). At this interface, the ChP receives a disproportionate share of cerebral blood flow^2,3^ via a rich network of fenestrated capillaries^1^ and mediates a robust two-way exchange of blood and CSF components. In turn, the ChP produces the CSF (400-600 mL per day in humans)^1^, which equilibrates with interstitial fluid of the brain and spinal cord across permeable linings (pia and ependyma) and via glymphatics^4^. CSF then returns to the peripheral circulation via arachnoid villi, granulations, and lymphatics in olfactory, dural, and basal regions^5^. These circulations and equilibria give the ChP and CSF access to CNS cells behind the blood-brain barrier, providing the rationale for CSF-based delivery of neurotherapeutics that cannot cross this barrier^6,7^.

ChP functions at the blood-CSF interface are largely executed by its epithelial cells (CPECs)^1^. Among their roles, CPECs produce the CSF, secrete most of the CSF proteome, gate and transport molecules and immune cells between the blood and CSF, detoxify the circulating fluids, and form the anatomical blood-CSF barrier. Despite this broad physiologic and therapeutic relevance, remarkably little is known about human CPECs. The myriad of secretory and absorptive functions attributed to CPECs are thought to be carried out by a single epithelial cell type^1^ that advances through developmental stages and asynchronous phases (“dark” and “light” cells^8,9^). Among CPECs, lineage differences relative to the embryonic roof plate^10,38,39^, and molecular differences based on ventricular origin have been described^11^, and recent studies from mice^12^ and from human ChP organoids^13^ suggest a mammalian CPEC subtype specialized for ciliogenesis. However, formal delineation of CPEC subtypes and their lineage relationships remain lacking.

To address these basic and applied gaps, we previously developed methods^14,15^ to generate derived CPECs (dCPECs) from mouse embryonic stem cells (ESCs) using an aggregate-organoid approach. (Note: “dCPEC” is used when referring specifically to the derived cells *in vitro*.) Together with studies from mouse explants^16^, the earlier derivations established fundamental developmental principles in mouse CPEC development, including BMP4 sufficiency as a CPEC morphogen^14,15^ and the temporal restriction of CPEC competency to pre-neurogenic neuroepithelial cells (NECs) rather than radial glia^14^. For *human* dCPECs, we used human ESCs and the neural rosette method^17^ to establish initial proof-of-concept^14^, but this method had limited utility due to its high complexity, inefficiency, and inconsistency.

In this study, we describe and validate a human dCPEC protocol with enhanced simplicity, efficiency, consistency, and scalability. Using this protocol, we then define multiple forms of CPEC diversity in humans. We first describe reciprocal multiciliated and Aβ uptake phenotypes that change over time, then use single cell RNA sequencing (scRNA-seq) across timepoints *in vitro* to define a human CPEC lineage tree with two bifurcations. Using independent derivations and perinatal tissues, we confirm and map the two bifurcations to midgestation and preterm periods in humans, then discuss the implications of this dynamic lineage to human brain development via two new models.

## RESULTS

### Simple, efficient, and accelerated human dCPEC generation from H1 ESCs

Our proof-of-concept method using human ESCs (H1 cells) yielded relatively few dCPECs (5-10%)^14^. To improve efficiency and simplicity, we optimized a feeder-free monolayer system involving the seeding of small ESC clumps on Matrigel followed only by media changes (Figs. 1A,B; S1A,B). Compared to an alternative high-density method, which also worked efficiently (Fig. S1C), the low-density seeding protocol provided ∼30X greater scalability and up to 30 million dCPECs from 2 million starting ESCs.

**FIGURE 1:**
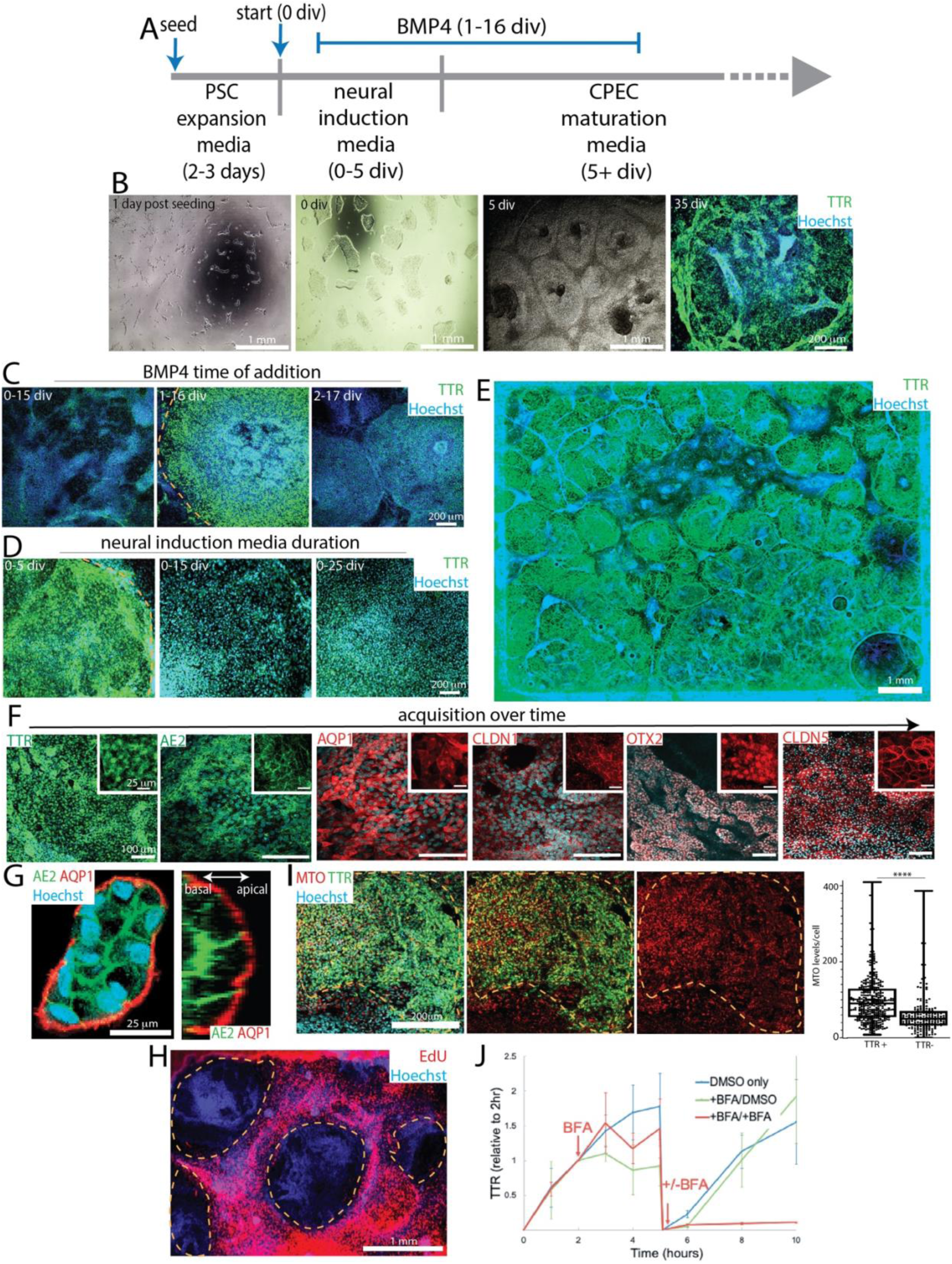
dCPEC derivation from H1 ESCs. Dashed lines demarcate dCPEC islands. **(A)** Protocol schematic. Small ESC clumps seeded at low density and subjected only to media changes. BMP4 treatment is 15 days starting at 1 div. **(B)** Derivation milestones (phase contrast and epifluorescent TTR ICC) include small low-density ESC colonies after seeding, colony growth to 150-200 um diameter at the start of derivation (0 div), confluent islands by 5 div, and 3D islands by 35 div. **(C)** BMP4 time-of-addition (epifluorescent TTR ICC; 20 div). 15-day BMP4 application starting at 1 div, but not 0 or 2 div, leads to strong dCPEC induction. **(D)** Neural induction media (NIM) duration (epifluorescent TTR ICC; 25 div). 5 days of NIM leads to better dCPEC differentiation than 15 or 25 days. **(E)** Example of an efficient derivation (epifluorescent TTR ICC; 45 div). Stitched images of one well in an 8-well chamber slide. **(F)** Marker acquisition order (confocal maximal projections; ICC): 35 div for TTR, AE2, AQP1, CLDN1; 50 div for OTX2; 70 div for CLDN5. **(G)** Apicobasal polarity (confocal and orthogonal views; ICC 45 div). Apical AQP1+ surfaces face the media, while AE2+ basolateral surfaces abut. **(H)** EdU incorporation (40 uM EdU; epifluorescent; 35 div). EdU application from 31-35 div leads to robust labeling of proliferating non-dCPECs (red), but not dCPEC islands. **(I)** Mitochondrial activity (confocal maximal projections of 150 nm MTO with TTR-ZO1 ICC; 35 div, ZO1 not shown; box plots designating medians, means “+”, quartiles and ranges) showing TTR+ dCPECs with significantly higher MTO levels (red) than non-dCPEC neighbors (t-test p<0.0001****; dCPEC n=367, non-dCPEC n=200). **(J)** TTR secretion into media following washouts (1 ug/mL BFA; ELISA; means +/- s.d.; 45 div). TTR increases in control wells (blue line) were reduced or blocked acutely by BFA at 2 hours (red and green lines), then increased in the absence of BFA after secondary washouts at 5 hours, indicating reversibility.

To accelerate dCPEC generation, “rapid” neural induction media (NIM; ThermoFisher) was introduced after ESC colonies reached 150-200 um in diameter (typically 2-3 days). As advertised, NIM led to ∼100% dual positivity by immunocytochemistry (ICC) for neural progenitor markers (NESTIN and SOX2) and negativity for the pluripotency marker OCT4 by seven days (Fig. S1D). Timing and duration of NIM and BMP4 (10 ng/mL) were then co-optimized. Consistent with the early and short window of NEC competency for mouse CPEC fate^14^, BMP4 addition one day in vitro (1 div) after NIM initiation, but not before (0 div) or after (2 div), resulted in substantial dCPEC induction (Fig. 1C). With BMP4 addition at 1 div, NIM for 5 days was most effective (Fig. 1D), after which the media was switched to CPEC media with BMP4. Like the mouse cells^14^, induction efficiency was similar over a range of BMP4 exposure times (10-30 days; Fig. S1E); 15-day BMP4 exposures were used for subsequent studies. Using this protocol, H1 ESCs formed dCPEC sheets and islands that routinely covered 40% or more surface area by 5-6 weeks (Figs. 1E; S1F) and could remain adherent and viable past one year. See Supplementary Information for detailed protocol information.

### dCPEC validations of canonical CPEC properties and functions

Like mouse dCPECs^14^, human dCPEC islands formed three dimensional (3D) folds^18^ and ridges over time and expressed established markers of several CPEC compartments by ICC (secretory apparatus/TTR^19^, apical membrane/AQP1^20^, basolateral membrane/AE2^21^, cilia/ARL13B^22^, nucleus/OTX2^23^, tight junctions/CLDN1 and CLDN5^24^) in a stereotypical order (Figs. 1F; S1G). The dCPECs also had a uniform apicobasal polarity, a well-known property of derived^14^ and *in vivo* CPECs^1^ (Fig. S1H), with apical AQP1+ membranes facing the media while basolateral AE2+ membranes faced Matrigel or abutted one another within 3D folds (Figs. 1G). Comparable derivation efficiencies were achieved with multiple iPSC lines (Fig. S1I-K; data not shown). As in mouse cells^25^, early cell cycle withdrawal was evident in human dCPECs, with relatively low EdU labeling after 21 div (Fig. S1L) and almost none after 31 div (Fig. 1H).

Consistent with the high mitochondrial content of CPECs *in vivo*^26^, dCPECs displayed high mitochondrial mass (ATPB; Fig. S1M) and activity (MitoTracker Orange; Fig. 1I) compared to neighboring cells. CPECs are also well known to produce and secrete TTR via the classical secretory pathway^27^. By 1-2 hours after washouts, TTR was readily detectable in culture media by ELISA (Fig. 1J) and blocked over a wide concentration range (0.1-10 ug/mL) of Brefeldin A (BFA), which inhibits the classical secretory pathway^28^. BFA blockade was also fully reversible (Fig. 1J), ruling out toxicity as an explanation for the TTR reductions.

### Dynamic multiciliated CPEC phenotypes *in vitro* and *in vivo*

CPECs are relatively unique among CNS cells in being multiciliated^29^. Even at low magnification, the cilia marker ARL13B readily distinguished monociliated non-dCPECs and neural rosettes^30^ (Fig. S2A) from the islands of dCPECs with multiple apical cilia per cell (Figs. 2A-B). Qualitative increases in cilia mass per dCPEC between 38 and 80 div (Figs. 2B; S2B-C) were confirmed using a metric based on ARL13B and ZO1 staining (Figs. 2C; S2D; see Methods). In addition, cilia occupied increasing percentages of dCPEC surface area over this time period (Fig. 2B,D).

**FIGURE 2:**
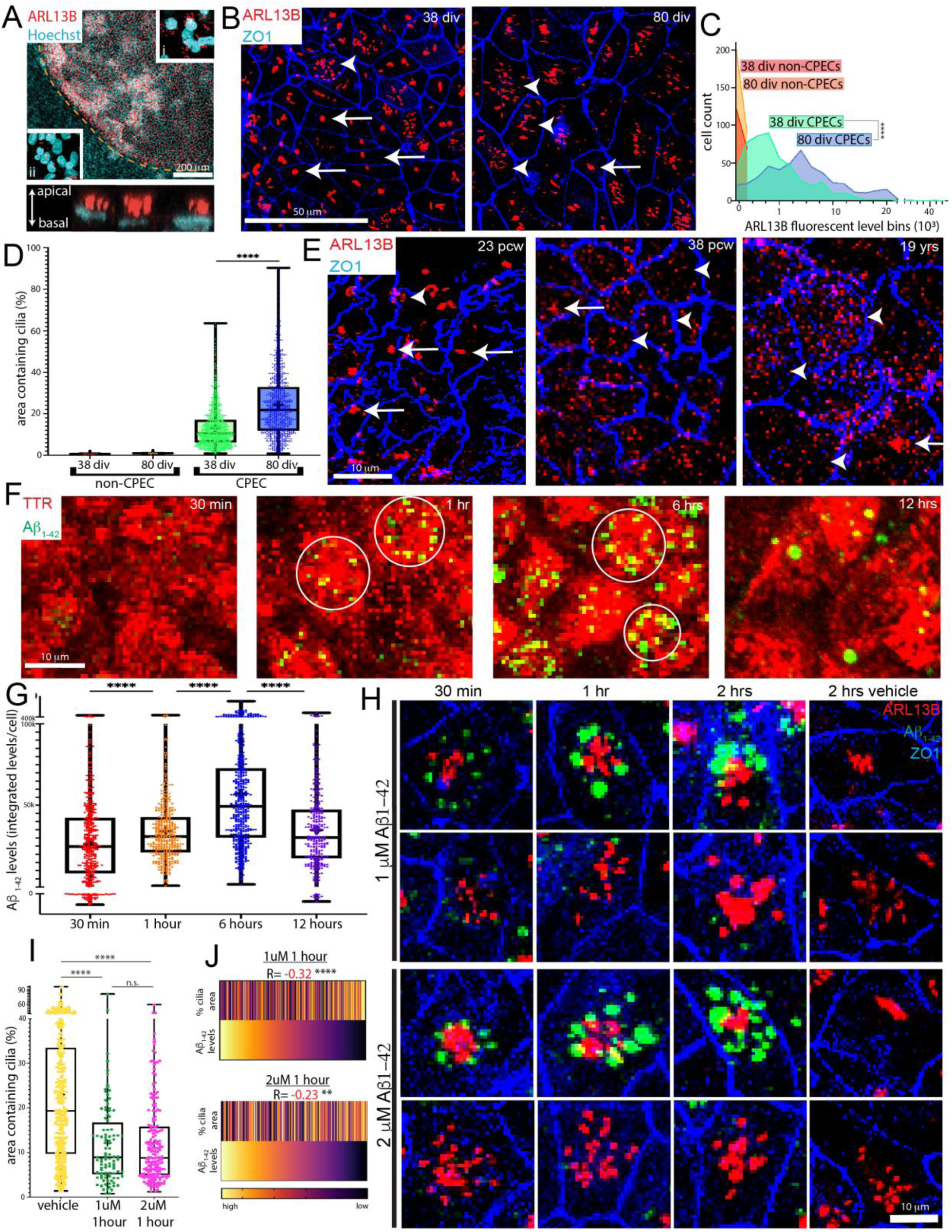
Dynamic and interacting multiciliated and Aβ_1-42_ uptake phenotypes. All images are confocal maximal projections. **(A)** Multiciliated phenotype (ARL13B ICC; 35 div); dashed line demarcates dCPEC island. Compared to monociliated cells (inset ii) outside the island, multiciliated dCPECs (inset i) are readily apparent even at low magnification. Orthogonal view (lower panel) demonstrates their apical localization. **(B-D)** dCPEC cilia mass increases and dispersion over time (ARL13B-ZO1 ICC; line graphs of cilia mass; box plots of surface area designating median, mean “+”, quartiles and ranges). dCPEC cilia are more clustered overall at 38 div than at 80 div (B). Cilia mass per dCPEC increases from 38 div to 80 div, while cilia mass in non-dCPECs remains low (one-way ANOVA p=0.0012, Bonferroni corrected t-test p<0.0001****; 38-div n=530, 80-div n=489) (C). dCPEC surface area containing cilia increases markedly between 38 and 80 div (one-way ANOVA p<0.0001, Bonferroni corrected t-test p<0.0001****; 38-div n=1189, 80-div n=923) (D). See Fig. S2D and Methods for metric details. **(E)** CPEC cilia dispersion *in vivo* (ARL13B-ZO1 whole mount ICC). Clustered cilia (arrows) are more frequent than dispersed cilia (arrowheads) at 23 pcw, but uncommon at 38 pcw and 19 years of age. **(F,G)** Aβ_1-42_ uptake and time course (fluorescent Aβ_1-42_ with TTR-ZO1 ICC; 64 div). Aβ_1-42_ uptake (green) into TTR+ dCPECs (red) often occurs in circular arrangements (circles in F), with uptake increasing and peaking by 6 hours, then decreasing by 12 hours (G; one-way ANOVA p<0.0001, Bonferroni corrected t-tests p<0.0001****; 30-min n=419, 1-hr n=415, 6-hr n=492; 12-hr n=300). See also Fig. S2F. **(H-J)** Aβ_1-42_ uptake and multicilia interaction (fluorescent Aβ_1-42_ with ARL13B-ZO1 ICC; 64 div). 1uM or 2uM Aβ_1-42_ uptake is robust by 1 or 2 hours in dCPECs with clustered cilia (H). After 1 hour, 1uM or 2uM Aβ_1-42_ results in dCPEC populations with more clustered cilia compared to vehicle controls (I; one-way ANOVA p<0.0001, Bonferroni corrected t-tests p<0.0001****; p=0.97^n.s.^; vehicle n=352, 1uM n=100, 2uM n=184). Rank-order heatmaps anchored by Aβ_1-42_ signal have significant negative correlations between Aβ_1-42_ levels and cilia clustering after 1 hour in 1uM or 2uM Aβ_1-42_ (J; Spearman correlations p<0.0001****, p<0.01**).

To assess cilia distributions *in vivo*, we evaluated whole mount preparations of human postmortem ChP using the same two antibodies (ARL13B and ZO1). Around midgestation (23 postconceptional weeks/pcw), multiciliated CPECs were abundant, and their cilia were generally clustered. Near term (38 pcw) and into early adulthood (19 years of age), CPECs remained multiciliated, but their cilia were widely dispersed (Figs. 2E; S2E). Thus, cilia clustering and dispersion *in vivo* paralleled that seen in dCPECs over time *in vitro*.

### Interacting multiciliated and Aβ uptake phenotypes

Robust uptake (absorption) is another fundamental property of CPECs^1^. Uptake of Aβ peptides associated with Alzheimer’s disease has been demonstrated for rat ChP^31^, but not yet for human CPECs. After treatment with fluorescently-tagged Aβ_1-42_ (AnaSpec), fluorescence in 64-div dCPECs was evident within 30 minutes, peaked by 6 hours, then decreased markedly by 24 hours (Figs. 2F,G; S2F) across a range of concentrations (500nM-4uM; data not shown). Aβ_1-42_ fluorescence was intracytoplasmic (Fig. S2G) within EEA1-labeled early endosomes, indicating rapid and robust Aβ_1-42_ uptake into the early endosomes of human dCPECs. EEA1 colocalization also decreased over time, suggesting Aβ_1-42_ transfer from early endosomes to other intracellular compartments (Fig. S2H,I).

Across concentrations and timepoints, intracellular Aβ_1-42_ appeared as variably-sized punctae and often in circular arrangements (Fig. 2F). ARL13B costaining revealed cilia at the center of these arrangements with Aβ-positive dCPECs having cilia that were generally more clustered (Fig. 2H). 1uM or 2uM Aβ_1-42_ application itself resulted in dCPECs having more clustered cilia than vehicle controls, with peak clustering one hour after application (Fig. 2I) and less by two hours (Fig. S2J). Moreover, Aβ_1-42_ signal intensities were inversely correlated with cilia dispersion (Fig. 2J). These findings suggest reciprocal interactions between cilia and Aβ_1-42_ uptake in human dCPECs.

### Direct dCPEC origin from tripotent neuroepithelial cells: Pseudotemporal scRNA-seq analysis

We then acquired four scRNA-seq datasets from 31 div (after apparent dCPEC cell cycle exit; Fig. 1H) to 75 div when dCPEC changes in morphology and immunocytochemical profile appeared to decelerate (Fig. 1F; data not shown). Timepoints were paired (31-46 and 55-75 div pairs; Fig. 3A) for culturing and processing through a 10x Genomics-Illumina pipeline to reduce batch effects, which we were unable to discern across the four datasets (Fig. S3A,B). Datasets were processed primarily through SoptSC, which performs unsupervised clustering, pseudotemporal ordering, and lineage inference in parallel^32^, as well as Seurat^33^.

**FIGURE 3:**
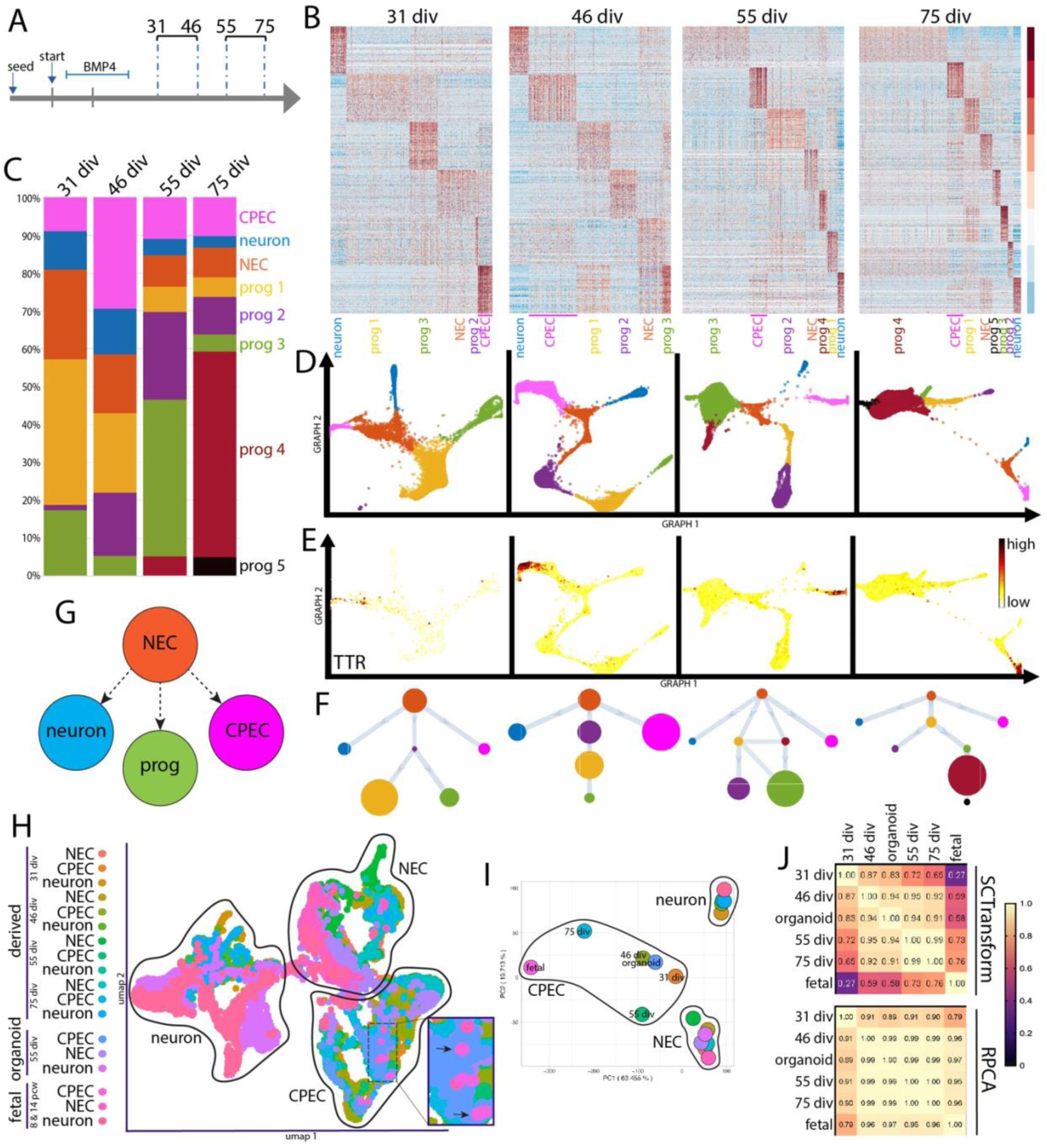
Direct dCPEC origin from tripotent NECs and comparison to human organoid and fetal CPECs. SoptSC analyses; see Fig. S3D-F for Seurat analyses. **(A)** Schematic of the four derivation timepoints and their pairings for culturing and scRNA-seq processing. **(B)** Heatmaps of top-300 DEGs per cluster. Cell types and color code designated below with dCPECs in pink. **(C)** Stacked bar graphs, color coded as in B. dCPEC percentage is highest at 46 div, then decreases as neural progenitors continue to proliferate and diversify. **(D)** GRAPH charts, color coded as in B. At all timepoints, NECs (light brown) have direct branches to dCPECs (pink) and neurons (blue). **(E)** GRAPH feature plots highlighting dCPECs (TTR expression). **(F)** Unsupervised SoptSC pseudotemporal lineages. NECs (light brown) have direct branches to dCPECs (pink) and neurons (blue) at all timepoints. NEC relationships to progenitors are more varied. **(G)** Lineage model for tripotent NECs giving rise to CPECs, neurons, and neural progenitors (“prog”). **(H)** UMAP of dCPECs, NECs, and neurons aggregated with organoid and fetal CPECs (SCTransform-corrected). CPECs from all sources form a cluster. Fetal CPECs cluster with older dCPECs (inset). **(I)** PCA of clusters, color coded as in H. The 75-div dCPEC cluster approaches the fetal CPEC cluster, while organoid CPECs display intermediate differentiation. **(J)** Pearson correlation tables using two batch correction methods. With SCTransform (top), CPECs display more differences. The dCPECs progress in an orderly temporal fashion towards the organoid, then fetal CPECs. RPCA correction (bottom) highlights CPEC similarities regardless of source. See Figs. S4C and D for complete correlation tables.

SoptSC analyses suggested six cell clusters at 31 div (Fig. S3C), which increased to eight by 75 div (Fig. 3B). Differentially-expressed genes (DEGs) and cell-type markers^11–14^ identified single clusters of dCPECs, neurons, and neuroepithelial cells (NECs) along with multiple clusters of neural progenitors – three at 31 and 46 div, four at 55 div, and five at 75 div (Fig. 3B,C). The same populations, with similar DEGs, were identified by Seurat (Fig. S3D-F). No non-neural cell clusters were identified in any dataset, which underscored the efficiency and consistency of neural induction and differentiation using the new protocol.

Consistent with NECs being stem progenitors, NEC fraction was highest at the earliest time point (31 div), while dCPEC and neuron fractions peaked at 46 div before declining (Fig. 3C). At 46 div, dCPECs comprised ∼30% of cells. By 55 and 75 div, dCPEC fractions decreased as additional classes of proliferating neural progenitors appeared, expanded, and contracted in an orderly fashion (Fig. 3C).

Pseudotemporal analyses of individual timepoints suggested three classes of NEC progeny. At all four timepoints, dCPECs arose directly from NECs as a direct and distinct end-branch (pink TTR-expressing cluster in Fig. 3D-F), as did neurons. Neural progenitors constituted the third progeny class, which had more varied relationships to each other and to NECs (Fig. 3D,F) consistent with their significant transcriptomic similarities overall. Among the three classes of NEC progeny (Fig. 3G), dCPECs had a consistently direct lineage relationship to NECs based on pseudotemporal ordering.

### Orderly dCPEC transcriptome changes compared to human fetal and organoid CPECs

The four scRNA-seq datasets were then compared to published human ChP organoid (55 div)^13^ and fetal CPECs (8 and 14 pcw)^34^. With batch correction by reciprocal principal component analysis (RPCA)^35^, CPECs from all sources clustered together and away from neurons and NECs by UMAP (Fig. S4A) and principal component analysis (PCA) (Fig. S4B). Batch correction using SCTransform^36^ increased the differences among CPEC clusters, which nonetheless continued to cluster and correlate well by UMAP and PCA (Figs. 3H,I). Pearson correlations between sources, using either batch correction method, supported dCPEC identity (Figs. 3J; S4C,D). These analyses also revealed increasing dCPEC-to-fetal CPEC correlations with increasing time *in vitro* (Figs. 3H-J; S4D), while the organoid CPECs displayed an intermediate differentiation profile (Fig. 3I,J) consistent with their intermediate derivation time (55 div)^13^.

### dCPEC subtypes and bifurcations: Pseudotemporal and time-series scRNA-seq analyses

Cells comprising the dCPEC lineage (dCPECs and NECs) were then re-run through SoptSC (Fig. S5A-B). At the two early timepoints (31 and 46 div), two dCPEC sub-clusters were evident (Fig. 4A-C). The sub-cluster with greater similarity to NECs was designated CPEC “type 1” (C1); the other sub-cluster was designated CPEC “type 2” (C2) (Figs. 4B; S5C). C2 fraction was highest at 31 div (∼40% of dCPECs compared to ∼10% at 46 div) (Fig. S5D), and C2 cells were present at the two later timepoints (Figs. 4D; S5F-H), but were too few in number to define as clusters computationally (see Methods).

**FIGURE 4:**
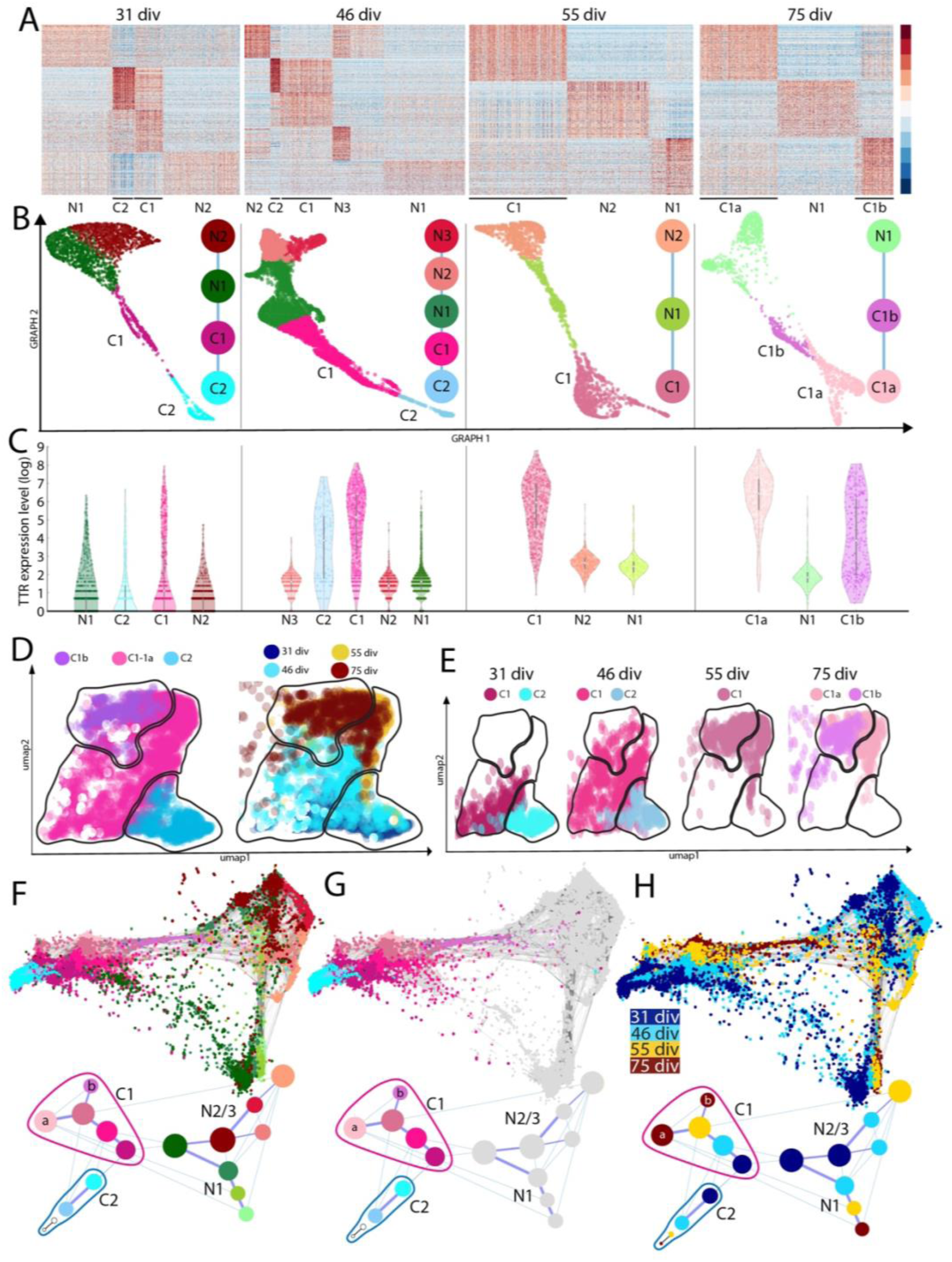
SoptSC and STITCH analyses across timepoints suggest a branching dCPEC lineage tree. **(A)** Heatmaps of the top-300 DEGs for dCPECs (“C”) and NECs (“N”) suggest two dCPEC subtypes at 3 of the 4 timepoints, but only one cluster at 55 div. **(B)** SoptSC GRAPH charts and lineage relationships coded by subtype. NECs are more similar to C1 than C2 cells. See Fig. S5C for pseudotemporal color coding. **(C)** Violin plots (with median and quartiles) of TTR expression, which increases in dCPECs over time with C1>C2 and C1a>C1b. **(D)** UMAPs of aggregated dCPECs, color coded and encircled for simplified subtype groupings (left) or timepoint (right). C2 cells (lower right) are mainly present early, but also at later timepoints, while C1 cells shift over time. **(E)** UMAPs of dCPECs from individual timepoints, color coded for dCPEC subtype as in B. As C2 cells contract, C1 cells adopt a “hybrid” C1a-C1b profile at 55 div before specifying into C1a and C1b subtypes by 75 div. **(F-H)** STITCH dot plots (upper) and lineage trees (lower), color coded for all subtypes (F), dCPEC subtypes only (G), or timepoint (H). The dCPECs cluster towards the left of the roughly-triangular dot plots, with NECs at the other two corners. Each corner has cells from all four timepoints (H). C1 (pink-purple) and C2 cells (blues) belong to distinct lineages from the outset. The C2 lineage displays transience, while the C1 lineage evolves and diversifies. Small 55-div and 75-div C2 “clusters” were manually added to the lineage trees to indicate their presence in low numbers at these timepoints.

Type 1 dCPECs were present at all four timepoints, but branched into two sub-clusters in the last 75-div timepoint (Figs. 4A-E). The sub-cluster with higher Pearson correlation (98%) to 55-div C1 cells was designated CPEC “type 1a” (C1a); the other sub-cluster (82%) was designated CPEC “type 1b” (C1b) (Fig. S5I). These correlations suggested the emergence of a distinct C1b subtype from the C1-C1a lineage. Similar sub-clusters and pseudotemporal relationships were obtained using Seurat (Fig. S5J-L).

For time-series analysis, dCPEC and NEC sub-clusters were analyzed using STITCH^37^. The roughly-triangular STITCH dot plots (Figs. 4F-H; S6A) suggested one broad dCPEC and two broad NEC groups. The dCPEC group contained dCPECs from all four timepoints (Figs. 4F-H; S6A). Additional STITCH analyses illustrated clear C1-C2 and C1a-C1b separations, supporting their designations as distinct subtypes (Fig. S6B). STITCH lineage trees were then formally constructed. At all timepoints, C2 cells were distinguished from the C1 lineage and NECs. The C1 lineage spanned all four timepoints, then bifurcated into C1a and C1b subtypes at the last timepoint (Fig. 4F-H, lower panels). Taken together, the SoptSC and STITCH analyses suggest two temporally-separated bifurcations within a branched dCPEC lineage tree.

We explored whether previously-described differences among CPECs could account for the dCPEC subtypes. CPECs in mice^38,39^ and humans^40^ have differing lineage relationships with the embryonic roof plate, but roof plate markers^38^ were not selectively associated with a dCPEC branch or subtype (Fig. S6C). Distinct anteroposterior (ventricular) CPEC identities have been described in mice^11,12^, but all dCPEC sub-clusters were mixtures of cells with lateral ventricle (LV), fourth ventricle (4V), or unspecified identities (Fig. S6E). Rapid initiation of dCPEC induction in our protocol (just one day after neural induction) is a likely contributor, although LV was favored over 4V identity (Fig. S6E), consistent with the “default” model of neural induction that favors anterior specification^41^. Regardless, neither roof plate lineage nor ventricular identity explained dCPEC branches or subtypes.

### Unique dCPEC ‘G0’ signature and direct origin from tripotent NECs: Cell cycle analyses

C1a-C1b lineage bifurcation *in vitro* occurred well after the apparent cell cycle withdrawal of dCPECs (Fig. 1H). Using Seurat^33^, which uses S, G2/M, and G0/G1 phase gene lists from mice^42^, dCPECs displayed G0/G1 or G1/G2M signatures (Fig. S7D); the G1/G2M dCPECs correspond to C2 cells due to the extensive overlap between ciliogenesis and G2/M genes^43^ (see next section). Individual dCPECs from all timepoints formed a cluster distinct from NECs, neurons, and neural progenitors by UMAP (Fig. S7D). When grouped by timepoint, dCPEC clusters separated from the other cell types by PCA (Fig. S7E). Interestingly, the three branches of NEC progeny (Fig. 3) were also clearly delineated by this cell cycle transcriptomic analysis alone (Fig. S7E-F).

We then used human cell cycle genes that distinguish a ‘neural’ G0 phase from G1^44^. The dCPECs from all stages had predominant G0 signatures, as did neurons (Figs. 5A,B; S7A,B), but dCPECs and neurons were clearly distinguished from one another, as well as from NECs and neural progenitors, by UMAP or PCA (Fig. 5A-D). Some C2 cells again displayed a G2M signature that nonetheless clustered with C1 cells (Fig. 5B,D). Notably, as with Seurat, human neural G0 analysis alone delineated the three progeny classes of tripotent NECs (Figs. 5C-E; S7C). Neural G0 analysis of the earliest dataset alone (31 div) delineated these classes (Fig. 5F).

**FIGURE 5:**
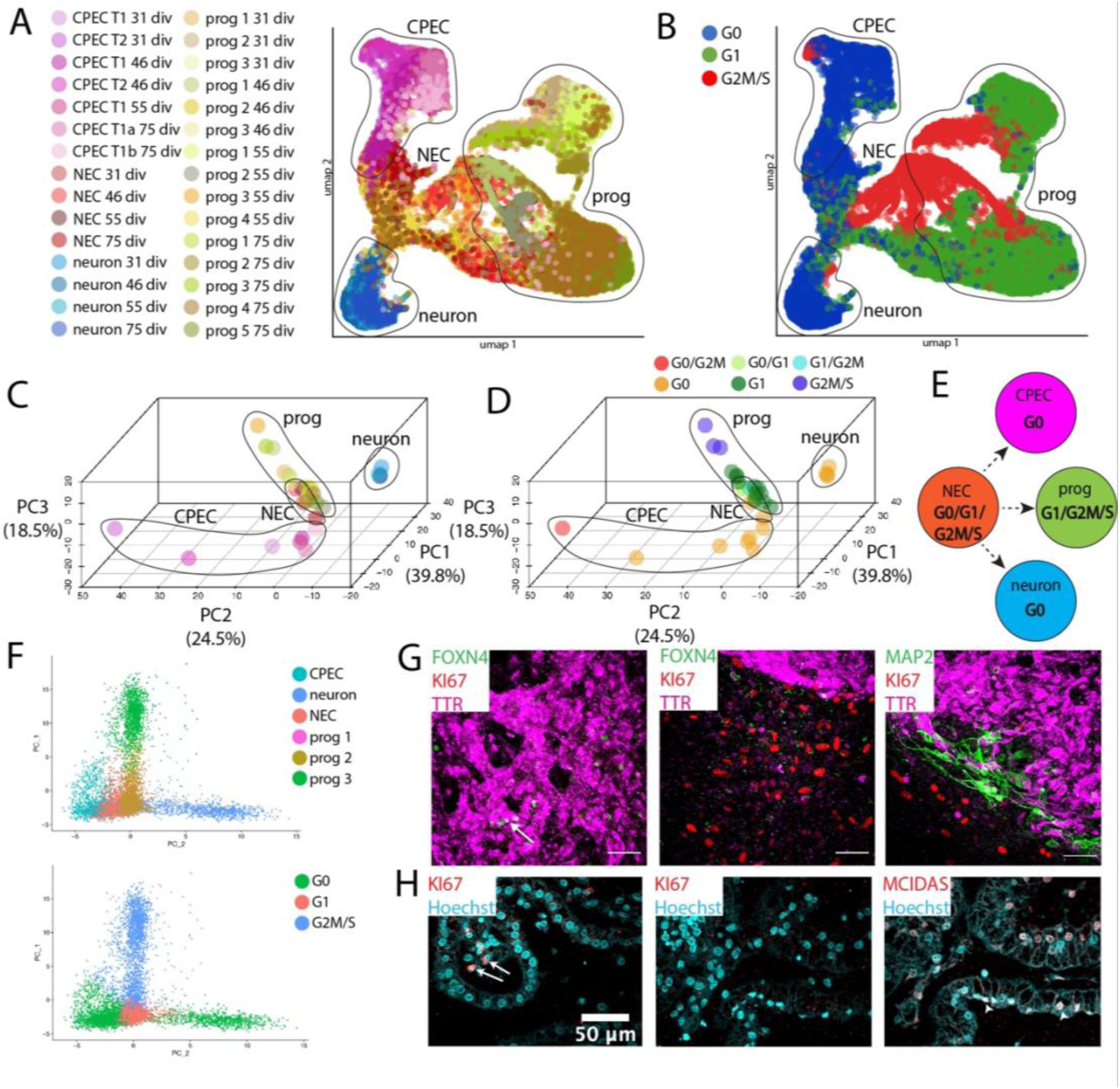
Cell cycle analyses highlight NEC tripotency and reveal a distinctive ‘neural G0’ dCPEC signature. Human **‘**neural G0’ transcriptomic analysis^44^ (A-F) with confocal maximal projections in derived cells (G) and 23 pcw tissue (H). **(A-B)** UMAP of all cells across timepoints, color coded for subtype and timepoint (A) or ‘neural G0’ phase (G0, G1, G2M/S), with major progeny classes encircled. The tripotent NEC progeny class structure is again evident (A). Across timepoints, dCPECs and neurons have predominant G0 signatures (blue), while NECs and progenitors are predominantly in G1 or G2M/S. **(C-D)** 3D ‘neural G0’ PCA plots of subtype clusters across timepoints, color coded for subtype as in A (C) or for predominant cell cycle phase(s) (D); see Fig. S7A,B for details. The three branches of NEC progeny are evident from neural G0 analysis alone (C). While dCPECs and neurons are principally G0 cells (orange), neural G0 analysis also distinguishes dCPECs from neurons (D). **(E)** Lineage model for tripotent NECs (Fig. 3G) with predominant cell cycle signatures added. **(F)** 2D ‘neural G0’ PCA scatterplot of all cells from the earliest timepoint (31 div), color coded for cell type (upper) or cell cycle phase (lower). NEC tripotency and dCPEC-neuron distinction are evident by 31 div. **(G)** KI67 status of dCPECs at an intermediate timepoint *in vitro* (TTR-FOXN4-MAP2-KI67 ICC; 49 div). KI67 (red) labels cells outside of dCPEC islands (middle panel), but not dCPECs (purple) or neurons (green in right panel). FOXN4+ C2 cells (arrow in left panel) also lack KI67 expression. **(H)** KI67 status of CPECs at midgestation *in vivo* (MCIDAS-KI67 IHC; 23 pcw); middle and right panels are immediately adjacent. Nuclear KI67 is present in some stromal cells (arrow in left panel), but not in CPECs, including MCIDAS+ C2 cells (arrowheads in right panel). See Fig. 6 for C2 marker studies using FOXN4 and MCIDAS.

To confirm their non-cycling status, we stained derived and perinatal CPECs for KI67, which labels cycling cells^45^. At a derivation timepoint (49 div) before C1a-C1b lineage bifurcation (Fig. 4H), KI67 clearly stained non-CPECs, but neither CPECs nor neurons (Figs. 5G; S7I). Similarly, in midgestation tissue (23 pcw) before C1a-C1b lineage bifurcation (see next section), KI67-positive cells could be seen in ChP stroma, but not in CPECs (Fig. 5H).

### Characterizations and confirmations of early type 1 and type 2 CPEC subtypes

All of the shared KEGG pathways^46^ and GO terms^47^ enriched in the top-300 DEGs of C1 and C2 cells at 31 div (when C2 cells were most abundant) were neurodegenerative diseases (Alzheimer, Huntington, Parkinson, and prion diseases; Fig. 6A). Selective C1 enrichments were associated with metabolism (e.g., metabolic pathways, non-alcoholic fatty liver disease) and energy (e.g., oxidative phosphorylation, thermogenesis), while C2 enrichments centered on ciliogenesis and cilia maintenance (Fig. 6A,B). Inspection of the top-10 individual DEGs at 31 and 46 div further highlighted the strong ciliogenesis signature of C2 cells (Fig. 6C). Expression of master ciliogenesis regulators^48^, particularly FOXN4^49^ and MCIDAS^50^, were collectively higher in C2 compared to C1 cells at all four timepoints (Figs. 6D,E; S8F). Secondary analysis of the human ChP organoids^13^ also identified a CPEC subpopulation that coexpress FOXN4 and MCIDAS (Fig. S6F).

**FIGURE 6:**
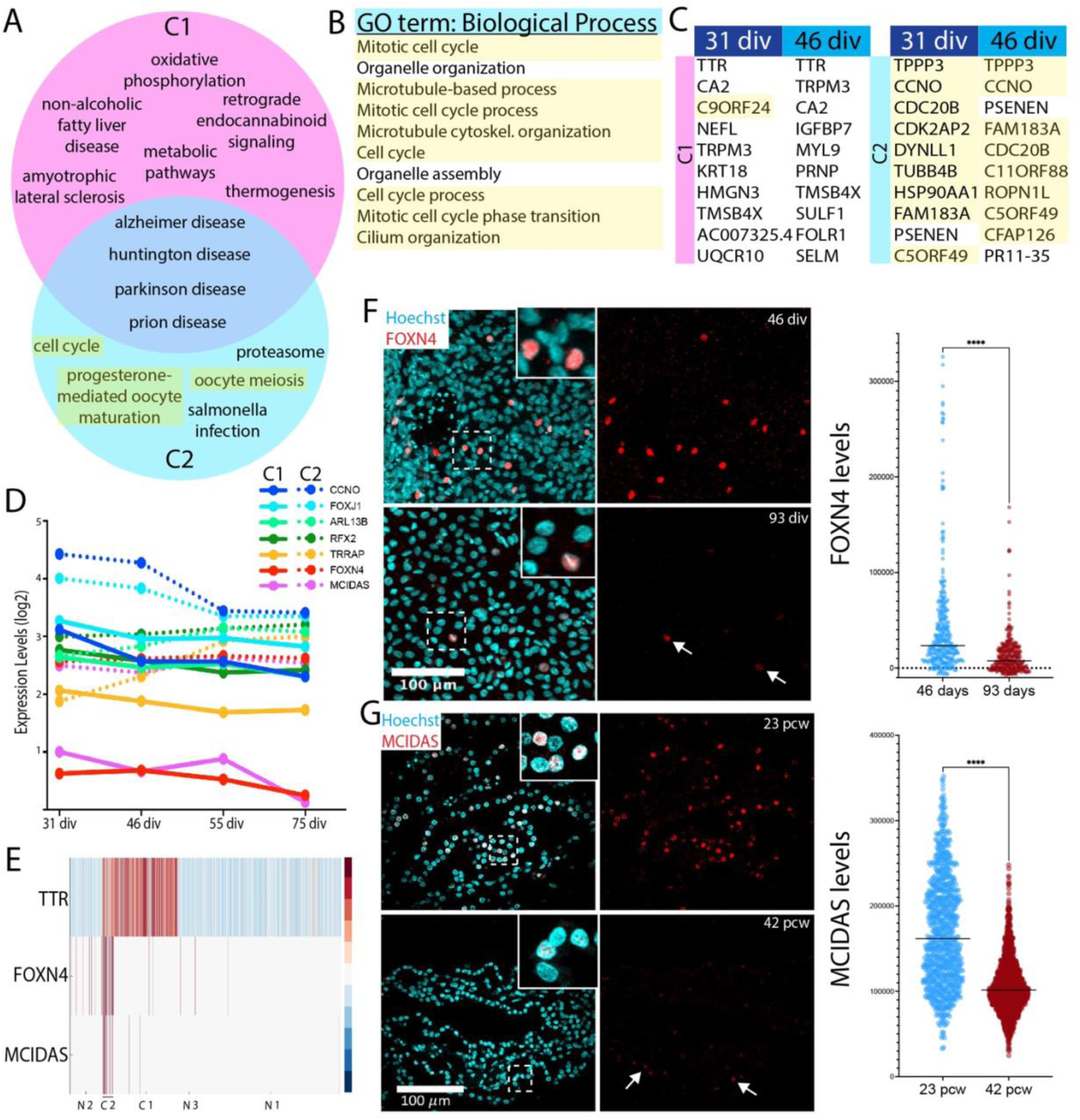
Early type 1 and type 2 dCPECs, with later type 2 contraction. **(A-B)** Venn diagram of all enriched KEGG pathways (p<0.05 with Bonferroni correction) (A) and top-10 GO terms for C2 cells (B) based on top-300 DEGs of C1 and C2 clusters at 31 div, with microtubule/ciliogenesis-associated terms highlighted. The C1 subtype (pink) is enriched for metabolic and energy-related pathways, while the C2 subtype (blue) is enriched for microtubule/ciliogenesis-associated terms. C1 and C2 subtypes share enrichment for neurodegenerative disease pathways. **(C)** Top-10 DEGs for C1 (left) and C2 subtypes (right) at 31 and 46 div, with microtubule/ciliogenesis-associated genes highlighted. C2 cells express ciliogenesis genes at particularly high levels. **(D)** Gene expression levels (log_2_) of seven master regulators of ciliogenesis^48^. C2 cells (dashed lines) express higher levels than C1 cells (solid lines), particularly FOXN4 (red) and MCIDAS (purple). **(E)** Heatmap of NEC and dCPEC subtypes at 46 div. FOXN4 and MCIDAS distinguish C2 from C1 cells and NECs. **(F)** C2 contraction *in vitro* (FOXN4 ICC maximal projections and violin plots with medians; 46 and 93 div). FOXN4 fractional positivity and expression levels decrease between 46 and 93 div (t-test p<0.0001****; n=300 each), although C2 cells are detectable at 93 div (arrows). **(G)** C2 contraction *in vivo* (MCIDAS IHC maximal projections and violin plots with medians; 23 and 42 pcw). Like FOXN4 *in vitro*, MCIDAS fractional positivity and expression levels decrease between 23 and 42 pcw (t-test p<0.0001****; 23 pcw n=799, 42 pcw n=1504), although C2 cells are detectable at 42 pcw (arrows).

FOXN4 and MCIDAS protein expression were then examined in independent derivations. Both were expressed at high (Fig. S8A) and highly-correlated levels (Fig. S8C) in early dCPECs (31 div) with FOXN4 demonstrating particularly discrete and mosaic expression (Figs. 6F; S8A,B). Over time, FOXN4-expressing C2 cells remained detectable, but significantly reduced in fractional positivity and expression level (Fig. 6F). Likewise, MCIDAS expression levels decreased over derivation time (Fig. S8A).

We then examined perinatal ChP tissues using the same two antibodies (FOXN4 and MCIDAS). Around midgestation (23 pcw), MCIDAS levels correlated positively with those of FOXN4 (Fig. S8D) with MCIDAS demonstrating particularly discrete and mosaic expression in a substantial CPEC fraction (Fig. 6G). Near term (42 pcw), MCIDAS-expressing C2 cells were detectable, but significantly reduced in fractional positivity and expression level (Fig. 6G). Similarly, FOXN4 expression levels decreased from midgestation to term (Fig. S8E). Consistent with the scRNA-seq and ICC findings *in vitro*, the perinatal tissues demonstrate higher type 2 CPEC prevalence at midgestation, which decreases significantly by term.

### Characterizations and confirmations of late type 1a and type 1b CPEC subtypes

Based on the top-300 C1a and C1b DEGs, C1b cells were uniquely enriched for endocytotic, stress response, and catabolic KEGG pathways (Fig. 7A) and Gene Set Enrichment Analysis (GSEA)^51^ terms (Fig. 7B). Conversely, while also sharing several pathways with the earlier-stage C1 cells (Fig. 7A-C; Extended data), C1a cells were selectively enriched for several secretory, water transport, and anabolic pathways based on manually-curated GSEA gene lists and GSEA plots (Fig. S9A; see Supplementary Information). GSEA analyses also revealed additional enrichments in C1b cells, such as immune cell signaling interactions and steroid hormone biosynthesis (Fig. S9B). C1a and C1b cells continued to share neurodegenerative disease pathways (Fig. 7A). Pearson correlations between top C1a and C1b DEGs were similar between 31 and 46 div, increased for a subset at 55 div, then strongly diverged by 75 div (Fig. 7D,E). Thus, 55-div C1 cells had features of a primed or mixed intermediate in normal development and transdifferentiation^52^. Re-examinations of the human fetal^34^ and organoid datasets^13^ did not reveal C1a and C1b subtypes (Fig. S6F; data not shown) consistent with C1a-C1b emergence after the stages sampled in these studies.

**FIGURE 7:**
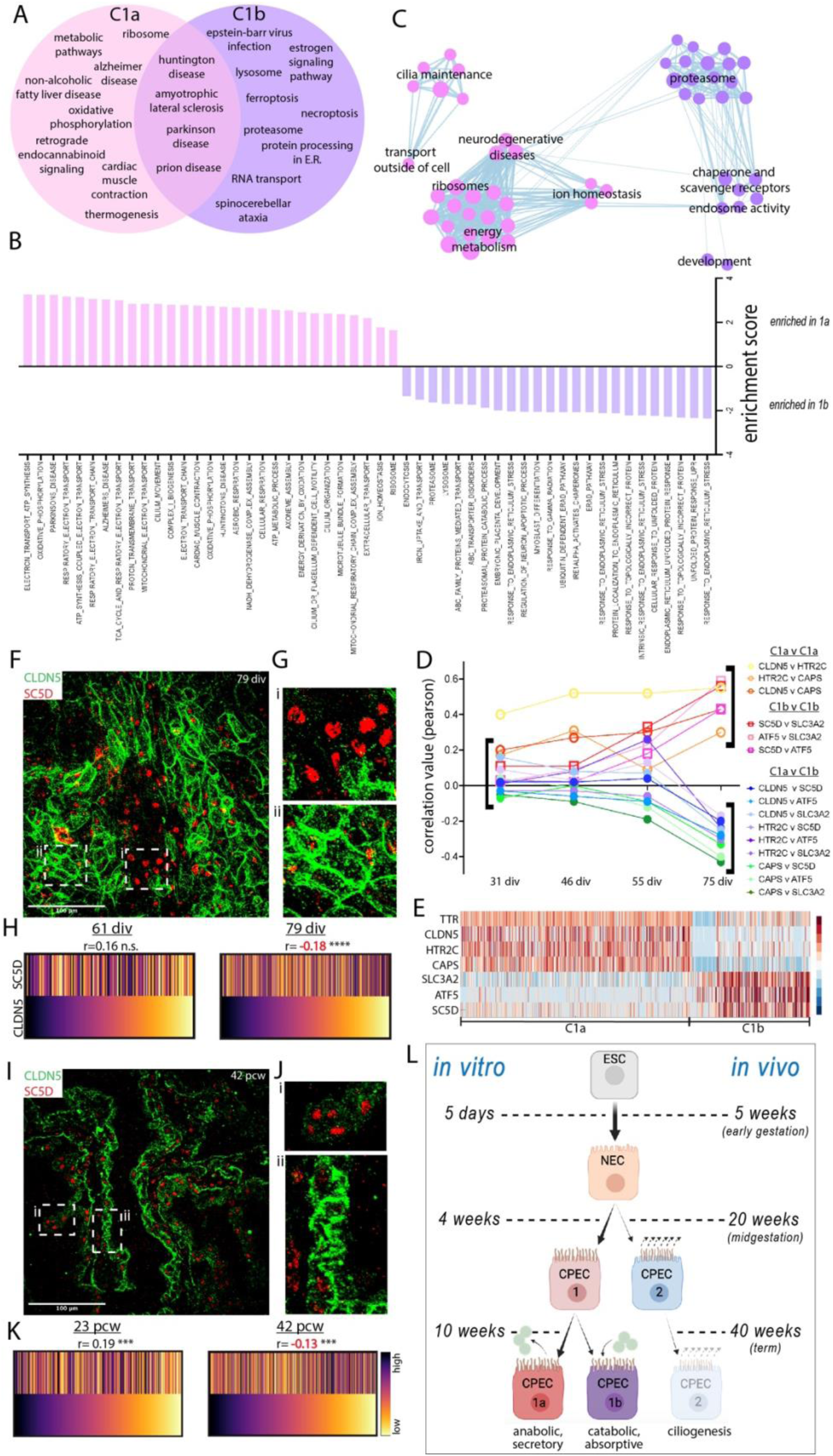
Late type 1a and type 1b dCPECs with prenatal CPEC lineage summary. **(A)** Venn diagram of enriched KEGG pathways (p<0.05 with Bonferroni correction) based on top-300 DEGs of C1a and C1b cells at 75 div. C1b cells (purple) are distinctively enriched for catabolic processes, while C1a cells (pink) share many metabolic and energy-associated pathways enriched in 31-div C1 cells (Fig. 6A). Directed GSEA analyses (Fig. S9A) demonstrate complementary enrichments of anabolic-secretory and catabolic-absorptive pathways in C1a and C1b cells, respectively. **(B)** Enriched GSEA terms in C1a (pink) and C1b cells (purple) using filtered genes (see Methods) expressed by >25% of cells. **(C)** Annotated GSEA theme plot illustrating C1a and C1b pathway enrichments. **(D)** Pearson correlations among (red-yellow) or between (green-violet) C1a and C1b marker genes. Before their uniform anti-correlation at 75 div, some C1a-C1b gene pairs display increased positive correlations at 55 div. **(E)** 75-div heatmap of C1a and C1b marker genes used in panels D-K and Fig. S9C-K. **(F-H)** C1a-C1b specification *in vitro* (CLDN5-SC5D ICC maximal projections with magnified insets and rank-order heatmaps with Spearman correlations; 61 and 79 div). Unlike CLDN5-SC5D coexpression at 61 div (Fig. S9F), discrete regions of CLDN5^hi^-SC5D^lo^ (C1a) and SC5D^hi^-CLDN5^lo^ (C1b) cells are evident by 79 div (F,G). A trend towards positive CLDN5-SC5D correlation at 61 div (p=0.2^n.s.^; n=150) becomes significantly negative by 79 div (p<0.0001****; n=584) (H). See also Fig. S9F-H. **(I-K)** C1a-C1b specification *in vivo* (CLDN5-SC5D IHC maximal projections with magnified insets and rank-order heatmaps with Spearman correlations; 23 and 42 pcw). Similar to *in vitro*, discrete regions of CLDN5^hi^-SC5D^lo^ (C1a) and SC5D^hi^-CLDN5^lo^ (C1b) cells are evident by the later stage (42 pcw). At 23 pcw, CLDN5-SC5D correlation is significantly positive (p<0.001***; n=323) before becoming significantly negative by 42 pcw (p<0.001***; n=812). See also Figure S9I-K. **(L)** Summary of the CPEC lineage tree with *in vitro* and *in vivo* timepoints.

Specification of C1a and C1b subtypes was then examined in paired derivations and human tissues. At 61 div, correlation between C1a (CLDN5) and C1b markers (SC5D) trended positively (Fig. S9D-F,H), then became significantly negative by 79 div (Figs. 7F-H; S9G,H). By 79 div, C1a and C1b subtypes also appeared clustered rather than randomly distributed (Fig. 7F,G). Similarly, in the human tissues, CLDN5 and SC5D expression were positively correlated at 23 pcw, then became negatively correlated by 42 pcw (Figs. 7I-K; S9I-K) and also appeared clustered (Figs. 7I,J; S9J). Together with the scRNA-seq analyses, these studies indicate a C1a-C1b lineage bifurcation that occurs between 61-75 div *in vitro* and between 23-42 pcw *in vivo*.

## DISCUSSION

In this report, we present an improved derivation of human CPECs, which differentiate with substantial cell autonomy and confirm several developmental principles from other species and systems^1,14^ (Figs. 1; S1). The human dCPECs also have dynamic multiciliated phenotypes that reciprocally interact with Aβ uptake (Figs. 2; S2), while pseudotemporal, time series, and cell cycle scRNA-seq analyses reveal their direct origin from tripotent NECs as well as substantial CPEC subtype diversity (Figs. 3-5; S3-7). Non-cycling type 1 (metabolic-energetic) and type 2 CPECs (ciliogenic) are present early *in vitro* and at midgestation *in vivo* (Figs. 6; S8). By term, type 2 CPECs contract (Figs. 6; S8) while type 1 CPECs bifurcate into type 1a (anabolic-secretory) and type 1b subtypes (catabolic-absorptive) (Figs. 7; S9). Transcriptomic comparisons within this human NEC-CPEC lineage model (Fig. 7L) suggest the ontogenetic and phylogenetic emergence of type 2 and type 1b subtypes from a primordial NEC-type 1-type 1a lineage.

### Robust human CPEC derivation with the new protocol

Molecular, cellular, functional, and transcriptomic findings established the identity and quality of dCPECs in the new protocol, which is more efficient, reproducible, scalable, and simple than our earlier methods^14,15^. In addition, dCPEC differentiation is relatively rapid and associated with phase-contrast milestones that are predictive and straightforward to assess (Fig. 1B; see Supplementary Information). Published CPEC derivation methods^13,53,54^ vary in their apicobasal topologies -i.e., whether apical or basolateral CPEC surfaces face the media. Using this protocol, dCPEC apical surfaces are media-facing (Figs. 1G; S1J); culture media therefore corresponds to the CSF compartment, which can be advantageous. While BMP4 suffices for human dCPEC induction, others have coapplied BMP4 with a WNT signaling activator (CHIR 99021) to induce CPEC fate^13,53,54^. In our hands, BMP4 alone induced dCPEC WNT expression (Fig. S6C), and CHIR 99021 provided no obvious additional benefit (data not shown).

### Developmental principles of human CPEC induction

The early BMP4 requirement (Fig. 1C) and direct NEC-CPEC lineage (Figs. 3-5) support NECs being the CPEC-competent progenitor rather than radial glia^14,55^, BMP4 sufficiency as a CPEC morphogen^14,15,16^, the cell-intrinsic nature of NEC responses to BMP4^16,56^, and the significant cell autonomy of CPEC differentiation^14,15^. At a minimum, absence of non-neural cells in the cultures (Figs. 3; S1D) indicates autonomy from mesenchymal, vascular, blood, and blood-borne influences, while the prominent 3D dCPEC folding (Figs. 1B,G; S1G,K) is consistent with ChP morphogenesis being driven by CPECs rather than mesenchyme^57^. Early dCPEC differentiation (Fig. 3) and cell cycle exit (Figs. 1H,5) also support CPECs being among the earliest differentiating cells in the human brain^38,40,58^.

*BMP4* and *MSX1*^56,16^ gene expression by dCPECs well after the cessation of exogenous BMP4 application (Fig. S6C), the autocrine/paracrine BMP signaling by dCPECs predicted by CellChat^59^ (Fig. S6D), and dCPEC island formation (Figs. 1; S1) suggest that human CPEC induction is homeogenetic (“like inducing like”), as postulated over a century ago^40^ and as described for the roof plate^60,61^ and telencephalic CPECs in mice^38,39^. BMPs, and BMP4 in particular, have been implicated in these homeogenetic phenomena, which expand the space and time over which morphogens can act during development^62^. By 55 div, type 1 dCPECs have a mixed identity (Figs. 4B,7D; S5H) typical of proliferative intermediates in normal development and transdifferentiation^52^ (Fig. 7D), but were non-cycling G0 cells (Figs. 1H,5). Thus, unlike the irreversible G0 of most neurons^63^, the G0 state of type 1 CPECs is presumably reversible.

### Dynamic and interacting Aβ uptake and multiciliated CPEC phenotypes

Initially-clustered CPEC cilia *in vitro* and *in vivo* dispersed over space and time (Figs. 2B-E; S2B,C,E). We found no literature precedent for such a phenomenon, but typically, clustered cilia are motile^64^, and CPECs in zebrafish^65^ and mice^66^ are known to have motile cilia. Interestingly, CPEC cilia motility in mice is transient^66^; cilia dispersal in human CPECs, along with type 2 CPEC contraction over time (see below), portend transience in humans as well. Conversely, the enhancement of cilia clustering by exogenous Aβ_1-42_ (Figs. 2I; S2J) raises the possibility that Aβ uptake promotes cilia motility. To our knowledge, this study provides the first demonstration of Aβ uptake by human CPECs, a potential Aβ clearance mechanism involved in Alzheimer’s disease initiation and pathogenesis^67^. Preferential Aβ uptake into early endosomes (Fig. S2H,I) surrounding clustered cilia (Fig. 2H) suggests that CPEC cilia serve as docking sites for uptake, as seen in other cell types^68,69^. Together with the enrichment of Alzheimer’s and other neurodegenerative disease pathways in all dCPEC subtypes (Figs. 6A,7A), the Aβ findings raise the possibility that CPECs and the ChP contribute to the developmental origins of Alzheimer’s disease^70,71^ and potentially other neurologic and neuropsychiatric disorders that manifest later in life^72,73^.

### Early type 1-type 2 subtypes and the multiciliated CPEC-to-ependymal transition

While dCPECs are generally multiciliated (Fig. 2A-E) and enriched for ciliogenesis genes compared to non-dCPECs (Fig. S8F), ciliogenesis enrichment further distinguishes type 2 cells (Fig. 6A-C). Type 2 CPECs correspond to the ciliogenesis-enriched CPECs described in developing mice^12^ and human ChP organoids^13^, with our study adding their early emergence, then contraction during prenatal human development (Figs. 4,6) and the cilia dynamics just discussed. Together with the aforementioned studies on CPEC cilia motility in zebrafish^65^ and mice^66^, our findings suggest a conserved CPEC-to-ependymal transition in multiciliated ventricular cells that promote CSF flow and mixing. Nonami et al.^66^ noted that mouse CPEC cilia motility precedes the emergence of ependymal cells^74^, whose cilia propel CSF^75^, but do not mature until weeks after birth^76,77^. Likewise, in humans, ependymal multiciliation and maturation are largely postnatal and continue into adolescence^78^, while CPEC differentiation^58^, CPEC cilia clustering (Figs. 2B-E; S2B,C,E), and type 2 cells (Figs. 4,6) are evident much earlier *in utero*. The prominent “bloom” of telencephalic ChP in cynomolgus monkeys and early first-trimester humans^58,62,79^ would further facilitate CPEC-mediated CSF flow and mixing before ependymal cells are able to.

### Late prenatal type 1a-type 1b subtypes and the preparation for birth

Type 1a-1b bifurcation does not occur until later *in vitro* or near term *in vivo* (Figs. 4,7), and this late prenatal timing suggests preparation for birth as the ontogenetic driver. Other than conception and death, birth is the most significant physiologic transition in human ontogeny due to the abrupt separation and loss of maternal, placental, and amniotic fluid supports for fetal energy flux (energy substrate delivery, usage, and waste clearance)^80,81^. Two well-known adaptations that support this dramatic transition in energy flux are the specification of surfactant-producing alveolar type II cells (AT2 or AEC2) in the lung and globin gene switching in the bone marrow. Both initiate during the late prenatal period with autonomy from the human birth process itself^82,83^ and directly impact the fluids for energy flux (air and blood) in neonates and adults - i.e., transport of oxygen, glucose, other energy sources, carbon dioxide, heat, and other forms of metabolic waste to and from tissues and the body.

Notably, CPECs also produce and directly regulate a fluid (CSF), albeit one that selectively subserves the brain. Hence, type 1a and 1b CPECs likely evolved to meet the extrauterine energy flux demands of the human brain, including the replacement of *in utero* functions provided by the placenta, which shares many of the same top enriched pathways as type 1b cells (e.g., waste removal, immune regulation, and steroid hormone production; Fig. S9B)^84^. While AT2 cells and globin gene switching are conserved among mammals^85,86^, type 1a and 1b CPECs have not been described previously. As mentioned, the human fetal^34^ and ChP organoid datasets^13^ lack these subtypes (Fig. S6F; data not shown), but were sampled before type 1a-1b specification occurs.

Regardless, while additional studies will be needed to define CPEC subtype phylogeny, *brain* phylogeny provides ample rationale for the ontogeny and phylogeny of unique CPEC subtypes in humans. Like AT2 cells and globin genes, fundamental CPEC roles center on energy flux and homeostasis needs, and these needs became massive in human brains. The absolute and relative expansion of hominin and human brains, especially their neocortices^87,88^, includes an exponential relative burst in *Homo sapiens*^87,89^ and associated changes in energy allocation that were just as dramatic. Despite representing only 2% of body weight, the adult human brain uses 20-25% of the energy budget compared to 8-10% in non-human primates and 3-5% in other mammals^90^, with the neonatal human brain expending a remarkable 60-65%^91^.

While CPEC subtype evolution to support the increased energy flux demands of human brains has not been described previously, molecular and physiological system adaptations have been. Notably, many of these rely on CPECs for their implementation in the brain. These adaptations include the thyroid hormone (TH) system, which also co-evolved with human encephalization^92^; the basal metabolic rate^93^ regulated by TH^94^; and the TH carrier and distributor system^95^, including TTR^96,97^, the most abundant CPEC product^11^ whose expression is enriched in type 1 and type 1a CPECs (Fig. 4C). These and other evolutionary adaptations would align with the preterm specification of specialized CPEC subtypes to support the uniquely energy-demanding human brain.

## Acknowledgments

We thank Melanie Oakes and the Genomics Research and Technology Hub (GRT Hub) for assistance with scRNA-seq, Vanessa Scarfone for assistance with FACS, the Sue and Bill Gross Stem Cell Research Center for access to core equipment and stem cell culture support, the Experimental Tissue Resource and autopsy service of the Department of Pathology & Laboratory Medicine for access to postmortem human tissue, and members of the Monuki lab for support and feedback on the work and manuscript. This work was supported by NIH R21 AG064640, NIH R21 MH109036, UCI SoM-SoBS pilot award, and Warren L. Bostick endowment (ESM); NIH R25 GM055246, NIH T32 NS082174, and Rose Hills Foundation (HM); NSF DMS1763272 and Simons Foundation 594598 (QN); and CIRM EDUC2-08383 (VE, OGJ).

## Data availability

Upon acceptance, RDS files for the sequencing data will be available through NCBI GEO. All codes used in this study were publicly available through bioconductor (<u>Bioconductor</u>) or through the institutions that published the platforms Seurat^33^ ^(^<u>Seurat - Guided Clustering Tutorial</u>), STITCH^37^ (<u>GitHub - wagnerde/STITCH: Assembles a graph manifold from time series single-cell</u> <u>RNAseq data</u>), and SoptSC^32^ (<u>GitHub - WangShuxiong/SoptSC</u>).

## Methods

### Pluripotent stem cell culture

H1 embryonic stem cells (ESCs; WiCell Research Institute, Madison, WI) and iPSCs (UCI ADRC iPSC bank) were cultured on growth factor-reduced Matrigel (Corning #354230) diluted in DMEM (Thermo Fisher #11995-065) and maintained in E8 media (Thermo Fisher #A1517001) at 37°C in 5% CO_2_ with daily full media changes. After reaching 80% confluence, PSCs were dissociated using ReLeSR following manufacturer protocol (StemCell Technologies #05872), passaged using E8 with 10uM Y-27632 (ROCK inhibitor; StemRD #Y-025). Y-27632 was removed after 12-24 hours. For quality controls, PSCs passed array CGH testing (Cell Line Genetics, Madison, WI) and pluripotency testing using a four-antibody kit (Thermo Fisher #A24881). Cell culture media were sterile filtered (Fisher Scientific #SCGP00525 and #0974063D), but no antibiotics or antimycotics were used per best practice guidelines^98^. Contamination was monitored by phase-contrast microscopy and mycoplasma detection kits (Thermo Fisher #M7006).

### CPEC derivation

Glass chamber slides (Millipore #PEZGS0816) were coated with growth factor-reduced Matrigel (0.085 mg/mL per chamber slide well). PSCs were dissociated, as above, and with trituration using a p1000 pipette tip to achieve clumps averaging 50 um in diameter. Cell solution was diluted to achieve 10 clumps/mm^2^ density before being distributed evenly in the wells and incubated at 37°C in 5% CO_2_. ROCK inhibitor was removed after 24 hours. Once PSC colonies reached ∼150 um in diameter (usually 2-3 days post-seeding), PSC media was fully replaced with Neural Induction Media (NIM; Thermo Fisher #A1647801), which marked the start of derivation (“N0”). 24 hours later (“N1”), media was replaced fully with NIM with 10 ng/mL BMP4 (R&D Systems #314-BP-050). On day N5, media was replaced fully with CPEC media composed of DMEM/F12 (Thermo Fisher #11330-032), NEAA (Thermo Fisher #07980), N2 (Thermo Fisher #17502048), and Heparin (StemCell Technologies #07980), supplemented with 10 ng/mL BMP4; day N5 became “C5”. After 15 days of BMP4 exposure (day C16), cultures were given daily half-media changes with CPEC media without BMP4 for the remaining time in culture. See Supplementary Information for additional details and optimizations of variables.

### EdU (5-ethynyl-2’-deoxyuridine) incorporation

The Click-iT EdU Alexa Fluor 555 kit (Thermo Fisher #C10338) was used per manufacturer protocol. Cultures were washed 2x with CPEC media, then given a full media change with 40 uM EdU in CPEC media, followed by daily half-media changes with EdU-CPEC media. At termination, cultures were aspirated, washed 1x with DPBS, then fixed with 4% PFA (FisherScientific #50-980-487) for 15 min at room temperature (RT). Click-iT reactions were performed following manufacturer protocol prior to immunostaining, Hoechst counterstaining, and coverslipping.

### Mitotracker Orange (MTO) labeling

MTO dye (Thermo Fisher #M7510) was reconstituted in DMSO (Thermo Fisher #D12345) per manufacturer protocol and used with minimal light exposure. Biological triplicate wells were gently washed 2x with CPEC media, then given full media changes with 150 nM MTO in CPEC media or vehicle in CPEC media. After 20 minutes at 37°C in 5% CO_2_, media was aspirated, cultures washed once with DPBS, then fixed with 4% PFA prior to immunostaining.

### Aβ uptake

FAM-labeled Aβ_1-42_ peptide (AnaSpec #AS-23526-01) was reconstituted following manufacturer protocol and used with minimal light exposure. Stocks were diluted to 500nM-4uM working solutions in CPEC media. Prior to treatment, cultures were gently washed 1x with DPBS, then CPEC-Aβ_1-42_ media was added to triplicate wells and incubated at 37°C in 5% CO_2_ for 30 min-48 hours. To minimize slide-to-slide variability, conditions were randomized across wells of different chamber slides. All conditions were stopped at the same time by aspiration, gentle washing 2x with DPBS, and fixation with 4% PFA for 20 min at RT.

### TTR ELISA with brefeldin A (BFA) inhibition

After 3x washes with warm CPEC media, 1ml of CPEC media was reapplied to each well of a 4-well chamber slide, then 20ul of conditioned CPEC media was longitudinally collected from individual wells over time (0-10 hrs starting with 1ml, ending with ∼820ul in each well). To inhibit secretion, BFA (Invitrogen #B7450) or vehicle (DMSO) was added directly to wells. To test later for BFA reversibility, media was aspirated, then cells were washed 3x before replacing with CPEC media with or without BFA. Longitudinally-collected media was assayed using a human TTR ELISA kit (AssayPro #EP3010-1) per manufacturer protocol and read on an HT microplate reader (Biotek Synergy). Background media-only values were subtracted, and concentrations were inferred from values normalized to the 2-hour timepoint using standard curves measured in parallel.

### Immunostaining

#### Antibody information

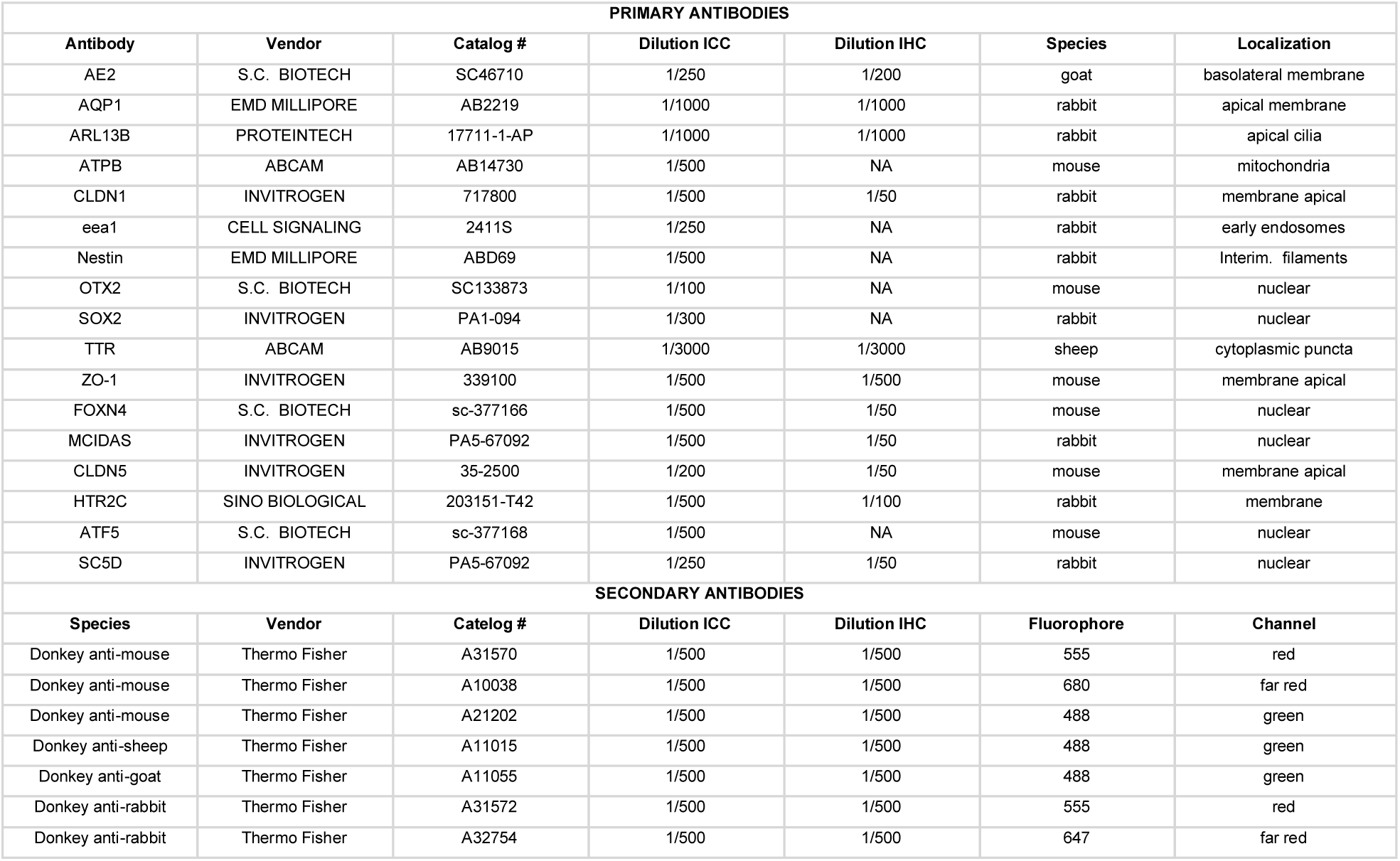

#### ICC of adherent cultures

Cultures were gently washed 2x with DPBS at RT, fixed with 4% PFA at RT for 20 min, then washed 3x with DPBS. Cultures were incubated in blocking solution (5% donkey serum (JacksonImmuno Research #017-000-121) in PBS + 0.03% Triton X-100 (MilliporeSigma #9036-19-5)) at RT for 1 hour, followed by incubation in primary antibody solution (primary antibody diluted in 1% donkey serum in PBS + 0.03% Triton) at 4C overnight. The next day, cultures were washed 3x followed by incubation in secondary antibody solution (secondary antibody diluted in 1% donkey serum in PBS + 0.03% Triton) at RT for 1 hour in the dark, washed 3x, then incubated in Hoechst solution (2ug/ml in PBS) (Thermo Fisher #H3570) for 5 min at RT in the dark, then washed 3x. The chamber slide apparatus was then removed per manufacturer protocol and slides were mounted in Fluoromount G (SouthernBiotech #0100-01) and coverslipped (Thermo Fisher #102460).

#### IHC of sectioned ChP

Paraffin-embedded human ChP tissue, fixed in 10% formalin at RT, were sectioned at 5um thickness by histology service (Experimental Tissue Resource). Sections were heated at 65C for 30 minutes, deparaffinized in xylene (Sigma-Aldrich #214736), then rehydrated through a descending alcohol series. Antigen retrieval using sodium citrate pH 6 (MilliporeSigma #6132-04-3) for 20 min in a vegetable steamer at 100C was followed by 1x ddH20 wash for 10 minutes at RT. Hydrophobic Pap-Pen barriers (Thermo Fisher #R3777) were drawn before fluorescent immunostaining and coverslipping, as described above.

### IHC of ChP whole mounts

Postmortem human ChP tissue from the lateral ventricle was obtained at autopsy, fixed and stored in 10% formalin at RT or 4C. Under stereo dissecting microscope, multiple small pieces (<1 mm diameter) were randomly picked and placed in a mesh basket (corning #CLS3478) in a 12-well plate. Free-floating immunostaining was performed as described above for sections using a rocker on low settings. Stained tissue pieces were placed on glass slides containing spacers (Invitrogen #S24735), then coverslipped.

### Image acquisition and modification

Phase contrast images of live cultures were acquired with an EVOS XL Core microscope (Advanced Microscopy Group). Fluorescent images were acquired with Olympus FV3000 confocal microscope, Nikon Eclipse E400 epifluorescent microscope with Nuance spectral deconvolution software, or Keyence BZ-X810 epifluorescent microscope for stitched images. All images were processed in ImageJ (v2.0.0). For all images compared qualitatively or quantitatively, identical image acquisition settings were used. Image adjustments in ImageJ were done in parallel and restricted to Brightness/Contrast.

### Image analysis and quantification

#### MitoTracker Orange

Confocal Z-stacks of cultures were converted to maximal projections. Membranous ZO1 signal was used to define TTR+ (CPEC) and TTR-negative (non-CPEC) ROIs, and total integrated MitoTracker Orange (MTO) signal was measured. Negative controls (no primary antibody) were imaged, then compartment-specific background signals were averaged and subtracted from experimental values. ROIs were drawn in blinded fashion prior to measuring integrated MTO values.

#### Cilia, Aβ uptake, and early endosomes

Confocal Z-stacks of cultures were converted into maximal projections. For cilia mass measurements, membranous ZO1 was used to define cell ROIs, then total integrated ARL13B (cilia) and/or fluorescent Aβ_1-42_ signals were measured. To assess cilia dispersion, outer contours of ARL13B signal were drawn and used as cilia ROIs. Cilia coverage was calculated by dividing cilia ROI area by cell ROI area. Negative controls were used for compartment-specific background subtraction, as above. For EEA1-Aβ colocalization measurements, the ImageJ co-loc2 tool was used. Due to inadequate signal using single cell ROIs, ROIs containing multiple TTR+ CPECs (5-10 cells) were drawn in blinded fashion before EEA1-Aβ colocalization measurements were taken.

#### CPEC subtype markers

Confocal Z-stacks of cultures and tissues were converted to maximal projections. For C2 markers (FOXN4 and MCIDAS), Hoechst channels were used to outline nuclear ROIs in ImageJ, which were then applied to the C2 marker channels to measure total fluorescent signal. For C1a and C1b markers, membranous CLDN5 signal was also used to outline cell ROIs. As above, ROIs were drawn in blinded fashion, and negative controls were used for compartment-specific background subtraction.

### scRNA-seq data acquisition and quality controls

Multiple dissociations methods were tested per 10x Chromium guidelines. The selected protocol involved dissociating with warm TrypLE (Thermo Fisher #12605010) for 25 minutes at 37C, resuspending in FACS buffer at RT (0.02% BSA in DPBS), then propidium iodine was added for 10 minutes (Thermo Fisher #P1304MP) before placing on ice. FACS for single live cells was performed by the UCI Flow Cytometry Core using a FACSAria II (BD Biosciences). Single cell capture with the 10x Chromium platform, cDNA library preparation, and cDNA QC analysis were performed by the UCI Genomics Research and Technology Hub (GRT Hub). Library generation was done with a V2 chemistry kit and an average capture of ∼10,000 barcoded cells, per user guide (CG00052 Rev B). The cDNA libraries were sequenced using the HiSeq 4000 platform (Illumina) to achieve an average depth of 50,000 reads per cell and 2,500 genes per cell. FASTQ files were processed and mapped to the reference genome GRCh38 through Cell Ranger. scRNA-seq datasets were then bioinformatically filtered to remove low quality cells and reads as previously described^12,32–34^. Briefly, genes expressed in <10 cells were excluded, and cells with 1000-6000 genes/cell and ≤12.5% mitochondrial gene expression were included.

See Supplementary Information for full details, including batch correction methods.

### scRNA-seq data analysis

#### SoptSC clustering and analysis

These were performed as described^32^. Briefly, the matrices of each time point were separately processed in MATLAB (v.R2019b). To determine cluster numbers, Eigengap heuristics were used, picking the lower boundary of the Eigengap, if supported by DEG heatmap. DEGs of known cell type markers were used to assign cell identity to each cluster.

#### Seurat clustering

Seurat^35^ v4.1 analysis was performed per online Seurat tutorial. Each time point was processed independently. Highly variable features with standard deviations ≥2.5 were used for PCA generation. Resolution ranged from 0.1-0.3, and DEG heatmaps were used to validate clustering and identity similar to SoptSC.

#### STITCH analysis

For STITCH^37^, highly variable genes were computed using Fano factors and ranking all genes by an above-Poisson noise statistic. Using default settings, top-2000 variable genes were filtered to include only genes whose single-cell transcript counts were minimally correlated (correlation coefficient >0.3) to at least one other variable gene. Gephi software was used for visualization. *Neural G0 analysis*

ScRNA-seq datasets were merged, then Seurat was used initially for cell cycle assignment. Using the same Neural G0 classifier^44^, either the mouse S and G2M gene lists from Seurat or the human G0, S, and G2M gene lists from Neural G0 were used. For Figure 5D, cell clusters were assigned their predominant cell cycle phase unless a second phase exceeded 30%, in which case a “dual” phase was designated (see legend for Fig. S7B).

#### CellChat analysis

For CellChat^32^, preliminary imputation using DrImpute was performed to account for dropouts, and the imputation output was used as input. The CellChat vignette was followed to create various communication and summary plots (https://github.com/jinworks/CellChat).

#### CPEC comparisons across datasets

Raw reads and metadata of published datasets^13,34^ were obtained from the UCSC Cell Browser. Datasets were processed through Seurat, as done by the originating groups, using the same quality control thresholds used for dCPECs. Cell types of interest (CPECs, neurons, NECs) were merged across datasets, followed by batch correction with either RPCA^35^ or SCTransform^36^ before clustering, Pearson correlations, and PCAs were performed in Seurat.

#### Gene enrichment and pathway analyses

Differential (top 300) and highly-expressed genes (top 10%) were used as input for pathway enrichment studies, then TopGO, KEGGprofile, and ReactomePA packages returned terms that reached statistical significance (p<0.05 with Bonferroni correction). GSEA enrichment analysis compared all genes expressed in ≥25% of cells, also returning terms with p<0.05 with Bonferroni correction. Heatmaps of manually-curated gene lists for specific CPEC subtype functions (Fig. S9A,B) were generated after mining the GSEA human molecular signature database (https://www.gsea-msigdb.org/gsea/msigdb).

#### Correlation and aggregate analysis

Evidence of early presence of the late emerging C1a and C1b subtypes were explored by correlating the expression of every pairwise combination (15 in total) of the top 3 C1a and C1b DEGs were generated across type 1 CPECs only in every time point with the ScatterFeature function to yield a pearson correlation value. Additionally, evidence for early presence of the late emerging subtypes (C1a and C1b) along with lingering presence of the early subtypes (C1 and C2) was explored by aggregating the CPEC and NEC subtypes across the four time points and subjecting it to clustering which generated 6 clusters, supported by DEG heatmap. UMAP graphs of the aggregate dataset were color coded and or split by time point or cluster of origin.

See Supplementary Information for additional details.

### Statistics, graphing, and AI attestation

For all studies, measurements used for statistical analysis were taken from distinct samples. Specific statistical tests are described in Figure Legends and Supplementary Figure Legends. All t-tests were unpaired and two-tailed, while all ANOVAs were one-way. Pearson correlations were used to compare transcriptomes datasets and fluorescent levels. For rank-order analyses, Spearman correlations were used. GraphPad Prism8 was used for graphing and statistical analysis of the ELISA, fluorescent measurements, and transcriptome correlations. Custom illustrations (Fig. 7L) were created with BioRender.com. All studies were conducted without the use of deep learning algorithms or platforms. Additionally, no form of AI was used for data analysis or writing this report.

### Reporting summary

Further information on research design is available in the Nature Portfolio Reporting Summary linked to this article.

## ETHICS DECLARATION

### Inclusion & Ethics

This study involved various local and regional researchers across every stage. The collaborators in this study qualify for authorship as outlined by Nature Neuroscience and are listed as authors. Their involvement and contribution were discussed prior to the start of the research. This work is locally relevant as determined by all collaborating researchers. These studies were carried out under the supervision and approval of local ethics committees (UCI IBC, IRB, and hSCRO). Postmortem ChP tissue studies were determined to be non-human subjects research by the UCI IRB. Research was conducted only after sufficient laboratory and safety training had been provided to all researchers. The research was neither restricted by the setting of the researchers nor resulted in stigmatization, incrimination, discrimination, or personal risk to participants. Local and regional research relevant to our study was addressed properly in the citations and the report.

## AUTHOR INFORMATION

### Contributions

H.M., B.A.J., and E.S.M. conceived the study. H.M., C.T., B.A.J., and E.S.M. contributed the theoretical basis for protocol optimizations, while H.M. and C.T. performed the optimizations. H.M. and Q.N. acquired the scRNA-seq data. H.M., S.W., Y.S., M.K.K., N.K., K.K., Q.N., and E.S.M. contributed to scRNA-seq analysis. H.M., P.S.R.F., B.T., and C.M. contributed to fluorescent imaging and quantification. H.M., V.E., B.A.J., and E.S.M. contributed to statistical analysis. H.M., M.N., and O.G.J. contributed to cell culture treatments. H.M., B.A.J., and E.S.M. contributed to experimental planning and data analysis. H.M. and E.S.M wrote the manuscript, and all authors approved the final manuscript.

### Corresponding author

Correspondence should be addressed to Edwin S. Monuki

---

**FIGURE S1:**
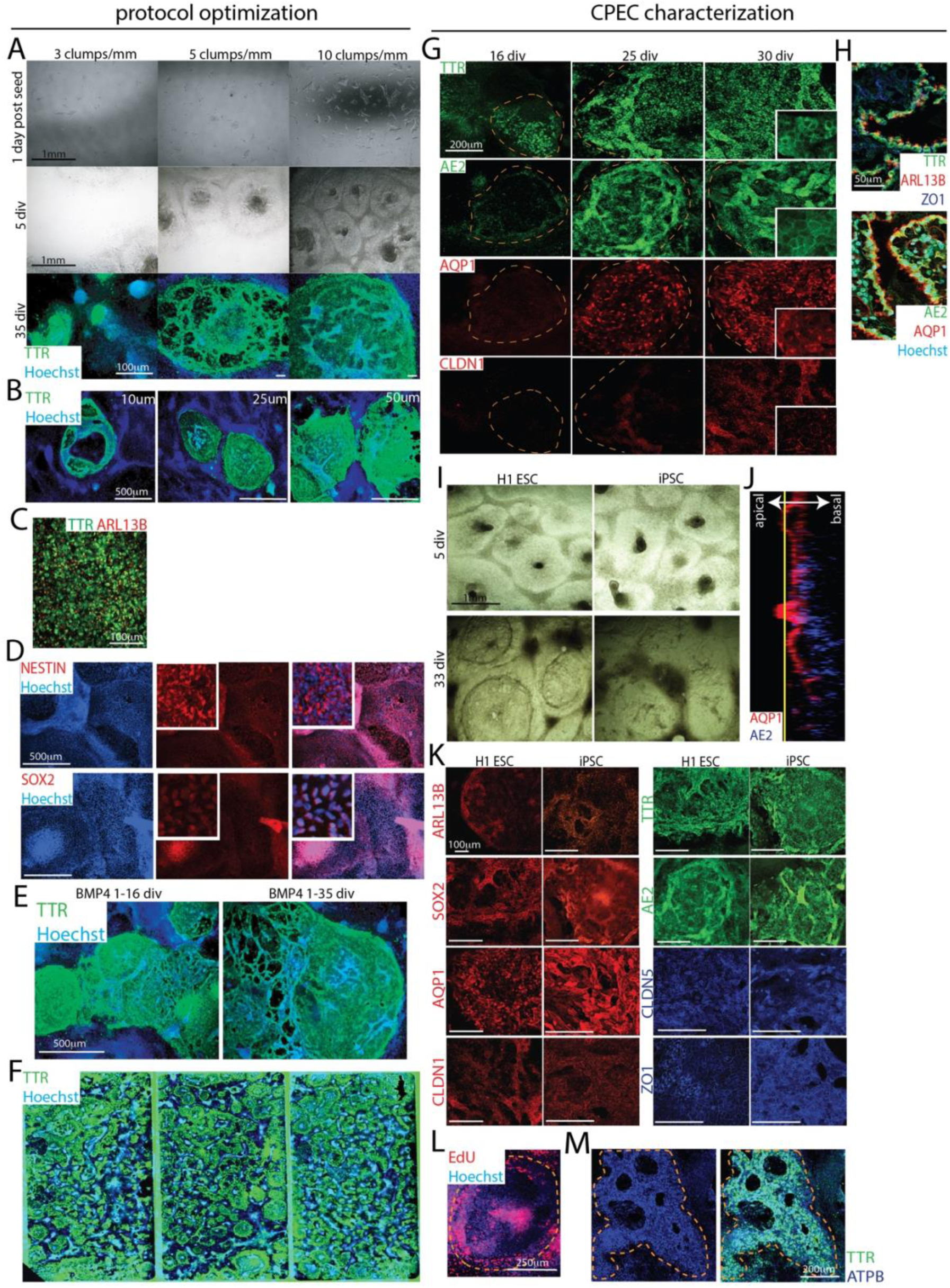
Derivation optimizations. Dashed lines demarcate dCPEC islands. Phase contrast (A,I), confocal maximal projections (H,J,M), and epifluorescent images. **(A)** Different ESC clump seeding densities (phase and TTR ICC). 10 clumps/mm^2^ yields more dCPEC islands than 3 or 5 clumps/mm^2^. **(B)** Different clump sizes at constant seeding density (TTR ICC, 35 div). Larger diameter clumps (50 um) yield more and larger dCPEC islands than 10 or 25 um clumps. **(C)** dCPEC induction using an alternative neural induction method (StemCell Technologies; TTR ICC, 35 div). **(D)** Uniform positivity for NESTIN and SOX2 after neural induction (ICC, 10 div). **(E)** BMP4 duration studies (TTR ICC, 35 div). No differences are apparent between 15- and 34-day applications. **(F)** Example of efficient and consistent dCPEC generation (TTR ICC, 35 div, stitched 4-well chamber slide). **(G)** Acquisition of dCPEC markers from 16-30 div (TTR, AE2, AQP1, and CLDN1 ICC). **(H)** Apicobasal polarity *in vivo* (39 pcw) with apical (ventricle-facing) ARL13B and AQP1, and basolateral AE2. **(I)** Similar phase-contrast appearance of H1 ESC and ADRC6 iPSC islands at key stages. **(J)** Uniform apicobasal polarity of ADRC6 iPSC-derived dCPECs (AQP1 and AE2 ICC, orthogonal view). **(K)** Similar CPEC marker expression in ESC- and ADRC6 iPSC-derived dCPECs (ICC, 35 div). **(L)** EdU application from 21-25 div leads to some EdU incorporation in 25-div dCPEC islands; see Fig. 1H for comparison (Click-iT reaction; ThermoFisher). **(M)** Prominent mitochondrial staining in TTR+ dCPECs compared to non-dCPECs (ATPB ICC, 35 div).

**FIGURE S2:**
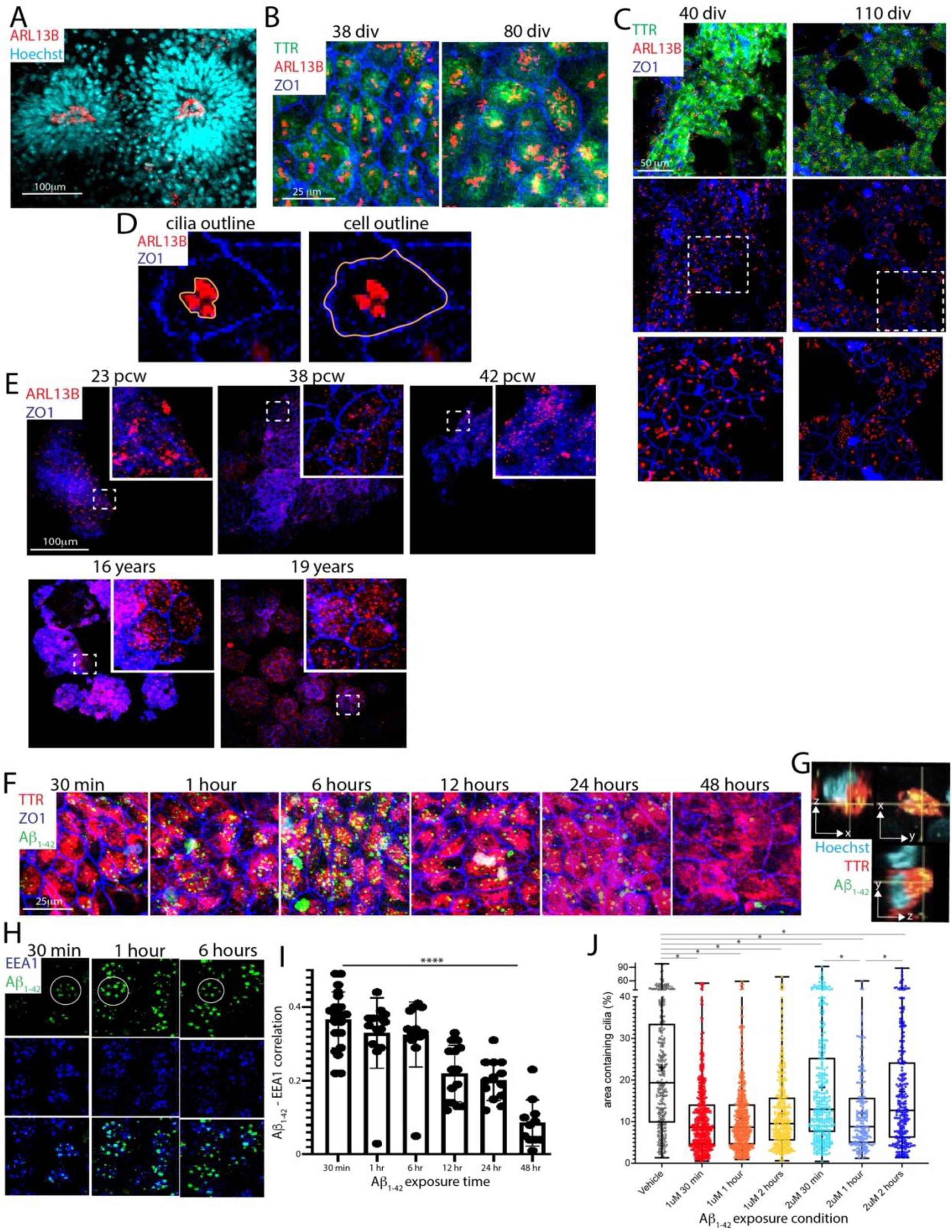
Multiciliated and Aβ_1-42_ uptake patterns in dCPECs. All images are confocal maximal projections. **(A)** ARL13B staining of neural rosettes, which is easily distinguished from dCPEC cilia staining. **(B)** Same field as Fig. 2B, but with TTR (green) to confirm dCPEC identity. **(C)** Another example of paired cultures displaying cilia clustering early (40 div) followed by dispersion (110 div). **(D)** Outlining approach for the cilia dispersion metric (ARL13B-ZO1 ICC; see Methods). **(E)** CPEC cilia staining in a more extensive *in vivo* series. ARL13B+ cilia remain dispersed in a 16- and 19-year old. **(F)** 1uM Aβ_1-42_ uptake by 64-div dCPECs over a more extensive time course. **(G)** Orthogonal views displaying intracytoplasmic colocalization of Aβ1-42 and TTR (64 div). **(H)** Aβ_1-42_ in EEA1+ early endosomes (64 div). Circular Aβ_1-42_ uptake patterns are apparent (circles). **(I)** Bar graph with data points (means +/- s.d.) showing decreased Aβ_1-42_- EEA1 colocalization over time (one-way ANOVA p<0.0001****; n=12 ROIs per timepoint). **(J)** Box plots (medians, means “+”, quartiles and ranges) of cilia clustering induced by 1uM or 2uM Aβ_1-42_ in 64-div dCPECs (ARL13B-ZO1 ICC). All Aβ-treated cultures had dCPECs with more clustered cilia than controls by one hour (one-way ANOVA p<0.0001, Bonferroni corrected t-tests p<0.05*; vehicle n=352, 1uM 30min n=531, 1uM 1hr n=476, 1uM 2hr n=354, 2uM 30min n=294, 2uM 1hr n=184, 2uM 2hr n=267).

**FIGURE S3:**
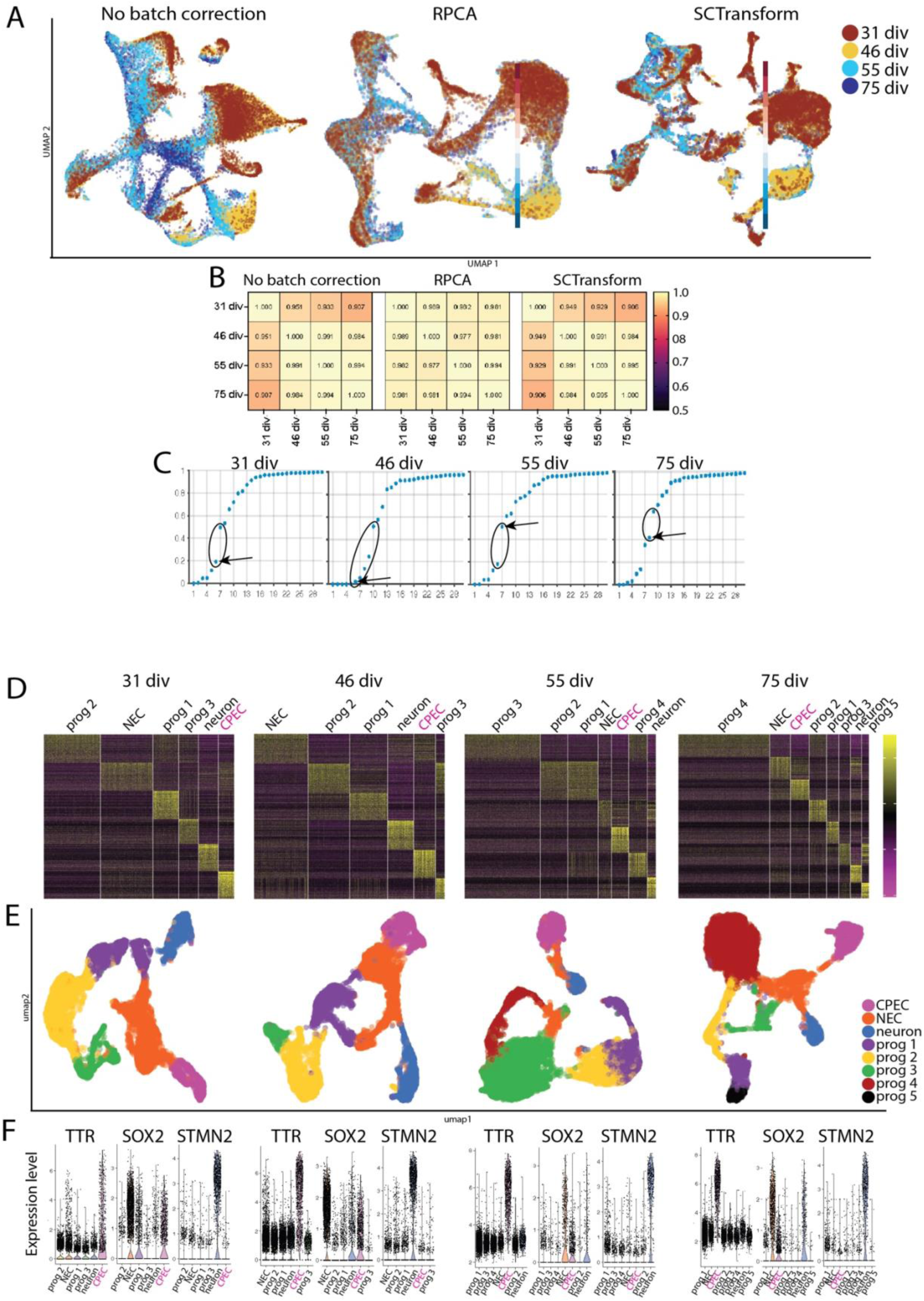
SoptSC quality controls and cluster number determinations (A-C) and Seurat analyses (D-F). (A) UMAPs of aggregated scRNA-seq data, color coded by timepoint, without and with RPCA or SCTransform batch correction. Clustering is discrete and spans timepoints in all cases. **(B)** Pearson correlations for dCPECs across timepoints, without and with RPCA or SCTransform batch correction. Correlations are generally high. Non-corrected and SCTransform-corrected correlations display perfect temporal ordering. **(C)** SoptSC cluster number determinations by Eigengap heuristic; ranges of bioinformatically-supported clusters are circled. Lowest cluster numbers were selected (arrows), except for 55 div, which yielded excellent DEG heatmaps using the higher number, as in D. **(D)** Heatmaps of top-100 DEGs support Eigengap heuristic-based cluster numbers. **(E)** Seurat UMAPs supporting SoptSC-determined cluster numbers and the direct origin of dCPECs (purple) from NECs (orange). **(F)** Violin plots of cell-type marker gene expression using Seurat-generated clusters. TTR highlights dCPECs, SOX2 highlights NECs, and STMN2 highlights neurons.

**FIGURE S4:**
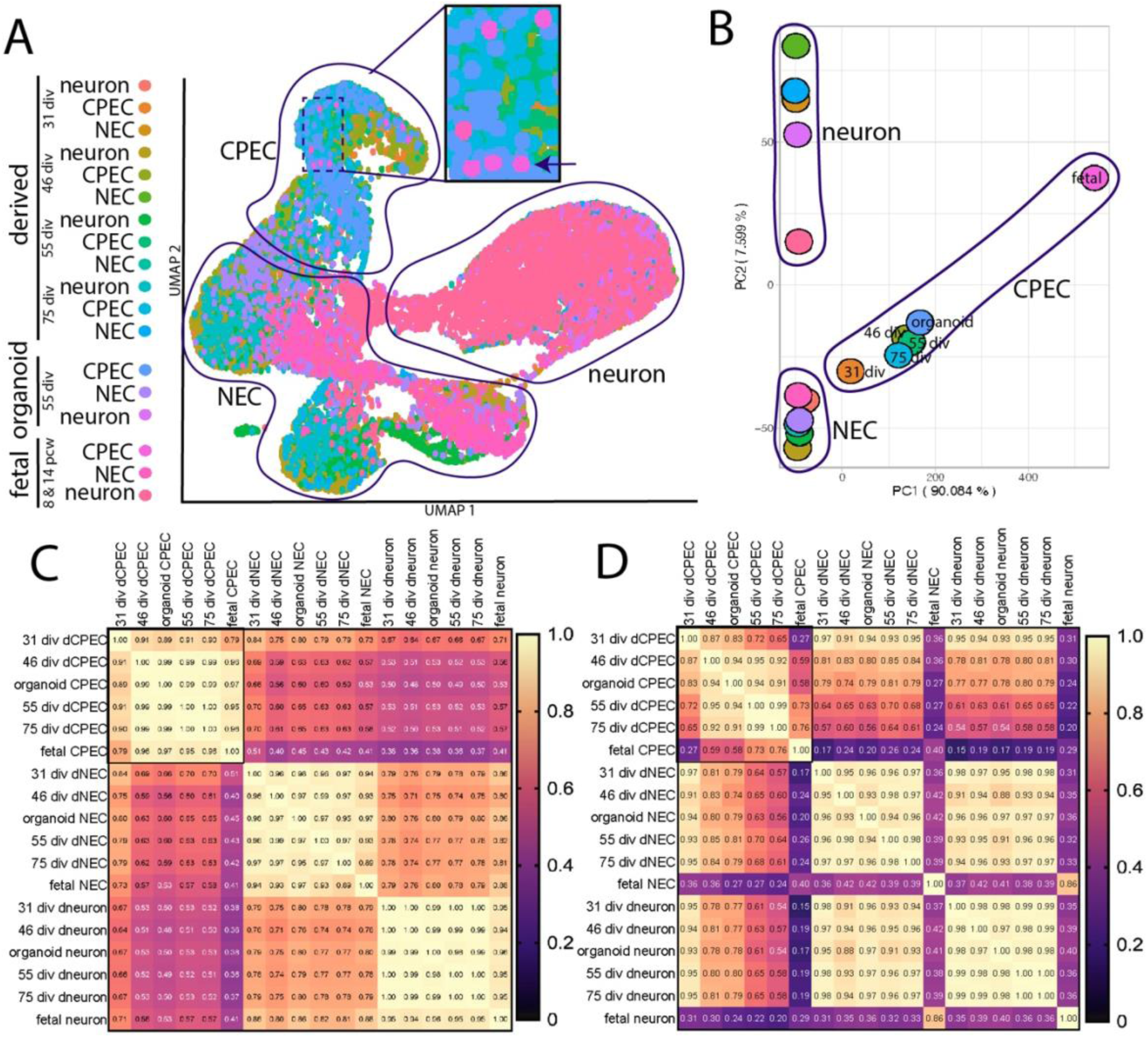
Additional dCPEC, organoid CPEC, and fetal CPEC comparisons. (A) RPCA-corrected UMAP of aggregated datasets. CPECs, NECs, and neurons from each dataset cluster well (outlined). Within the CPEC cluster, fetal CPECs (pink, arrow in inset) are more similar to older dCPECs (green-blue). **(B)** RPCA-corrected PCA of aggregated datasets, with CPECs, NECs, and neurons outlined. **(C)** RPCA-corrected Pearson correlation table for CPECs (boxed upper left), NECs, and neurons (lower right). Correlations between CPECs from the three datasets are high, as they are for NECs and neurons. **(D)** SCTransform-corrected Pearson correlation table, organized as in C. Correlations among the CPECs are generally lower, while NECs and neurons appear more similar.

**FIGURE S5:**
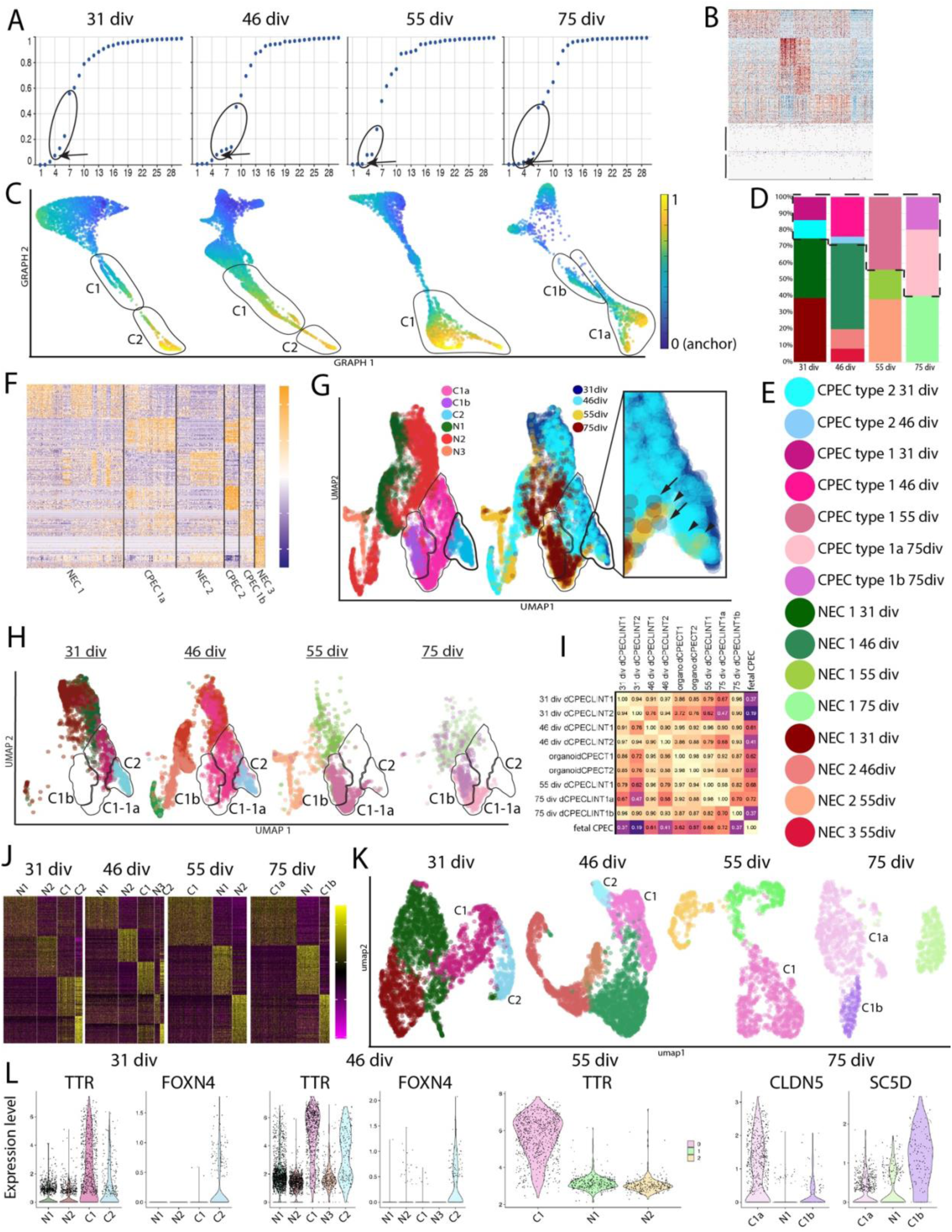
Additional SoptSC (A-I) and Seurat analyses (J-L) of dCPEC subtypes. (A) Eigengap heuristic for aggregated dCPECs and NECs. Ranges of bioinformatically-supported clusters are circled. The lowest cluster numbers were selected (arrows). **(B)** DEG heatmap example of overclustering. A 31-div heatmap for 6 clusters includes two without distinctive signatures (lower rows) and supporting 4 as the more appropriate choice. **(C)** Pseudotemporal color coding of SoptSC GRAPH charts shown in Fig. 4B, with NECs as anchor. NECs are more similar to C1 than C2 cells at 31 and 46 div. Both C1a and C1b cells have similarities to NECs. **(D-E)** Stacked bar graphs of dCPEC and NEC subtypes, with color coding in E. Total dCPEC and C1 fractions increase over time compared to NECs. **(F)** Heatmap of top-100 dCPEC and NEC DEGs aggregated across timepoints, which suggests 3 dCPEC and 3 NEC clusters like the individual timepoint analyses in Fig. 4A. **(G)** UMAPs of aggregated dCPECs and NECs, color coded by subtype (left) or timepoint (right), with dCPEC subtype regions outlined. Some C2 cells at 55 div (arrowheads) and 75 div (arrows) are detected. **(H)** UMAPs per timepoint, as in Fig. 4E, plus NEC subtypes. Similar dCPEC patterns are seen, including C1 lineage adoption of a hybrid C1a-C1b profile at 55 div before bifurcating by 75 div. **(I)** SCTransform-corrected Pearson correlations for organoid CPECs, fetal CPECs, and dCPECs separated by subtype and timepoint. 75-div C1a dCPECs are more similar than C1b cells to 55-div C1 dCPECs. **(J)** Seurat heatmaps of top-100 DEGs, supporting dCPEC and NEC cluster numbers and designations based on SoptSC. **(K)** Seurat UMAPs supporting dCPEC and NEC cluster numbers and designations based on SoptSC. **(L)** Violin plots of dCPEC subtype marker expression in Seurat-generated clusters.

**FIGURE S6:**
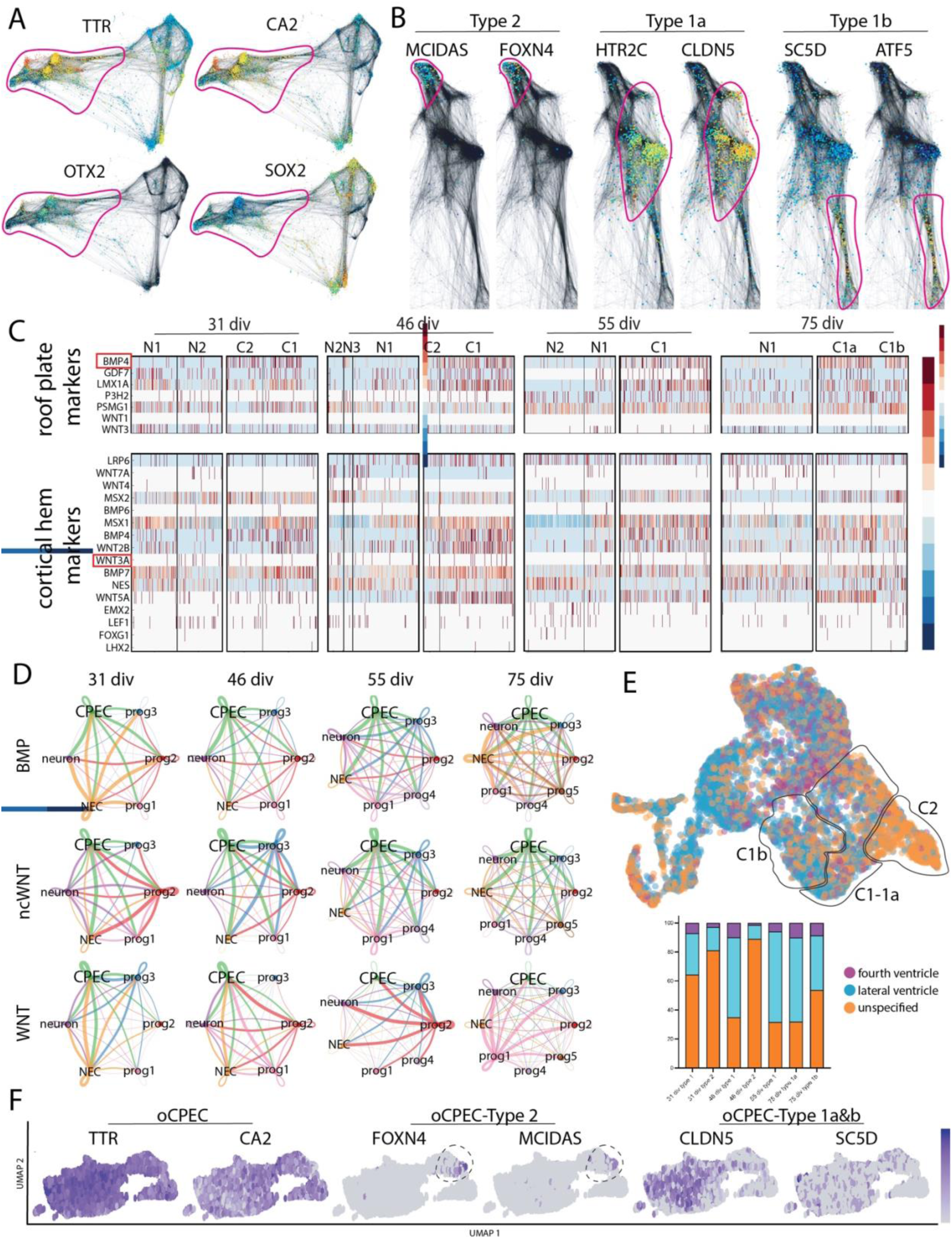
Additional bioinformatic analyses, including STITCH and CellChat. (A) STITCH dot plots of aggregated dCPECs and NECs, as in Fig. 4F-H, highlighting CPEC (TTR, CA2, OTX2) and NEC markers (SOX2); dCPEC regions outlined. **(B)** STITCH dot plots of aggregated dCPECs, highlighting type 2 (MCIDAS, FOXN4), type 1a (HTR2C, CLDN5), and type 1b markers (SC5D, ATF5); subtype regions outlined. **(C)** Heatmap of roof plate (top) and cortical hem (bottom) markers per timepoint. Patterns suggestive of unique relationships to dCPECs or dCPEC subtypes are not seen. Among BMPs, BMP4 and BMP7 are expressed by dCPECs, while WNT2B and WNT5A are expressed among WNTs. **(D)** CellChat analyses of major cell types for BMP, non-canonical WNT (ncWNT), and WNT signaling per timepoint. Predicted autocrine dCPEC signaling (green loops) and dCPEC signaling to other cell types (green curves) are prominent. **(E)** Ventricular (regional) specification of aggregated dCPECs and NECs using mouse genes^11^. Lateral (blue) exceeds 4th ventricular identities (purple), but identities are mixed, include substantial unspecified fractions, and do not distinguish dCPEC subtypes. **(F)** Organoid CPEC^13^ (oCPEC) expression of dCPEC subtype markers. Organoid CPECs express pan-CPEC markers (left) and include a C2 subpopulation (circled in middle), but C1a (CLDN5^hi^SC5D^lo^) and C1b (SC5D^hi^CLDN5^lo^) subpopulations are not distinguished.

**FIGURE S7:**
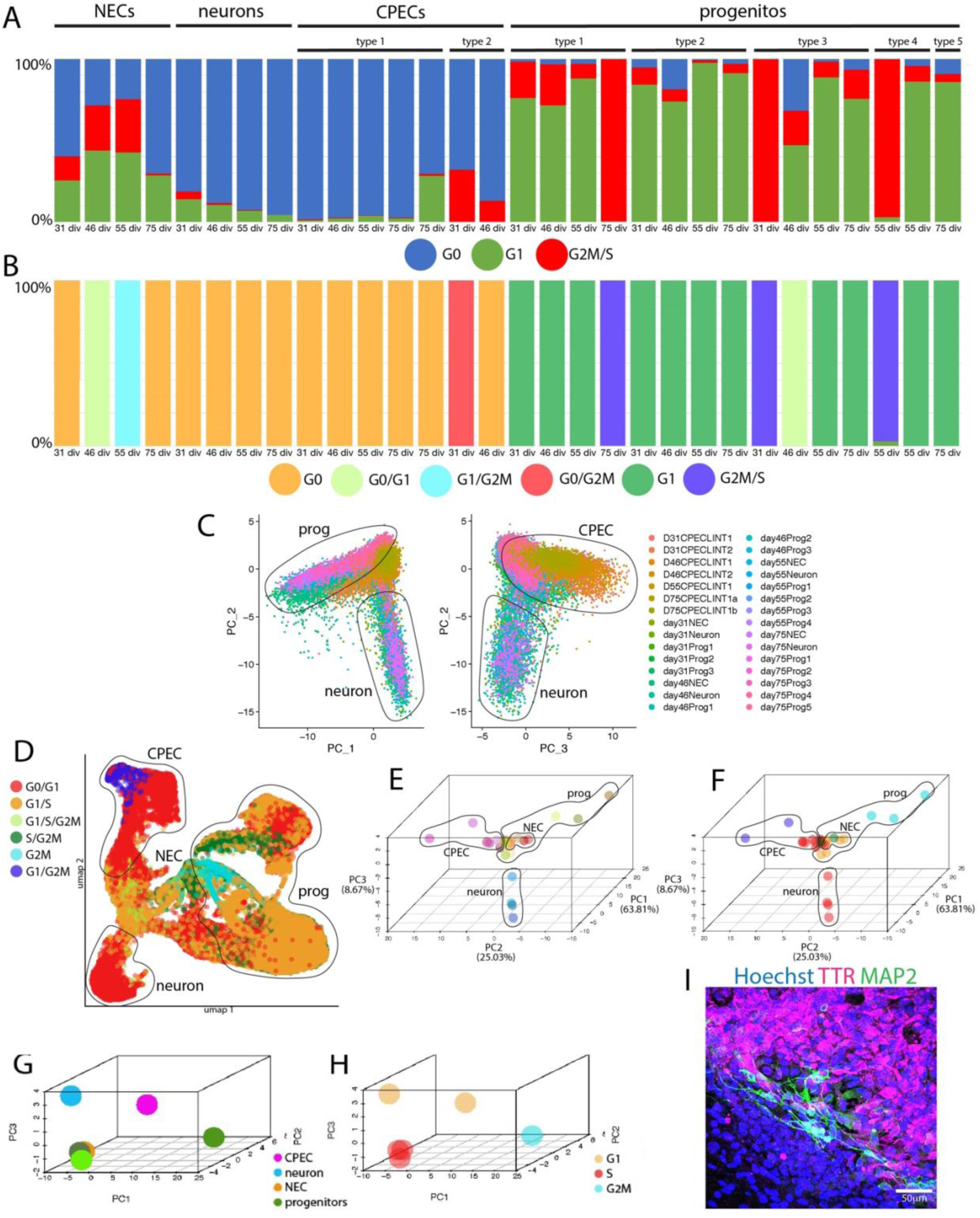
Additional cell cycle analyses using ‘neural G0’ (A-C) and Seurat (D-H). (A) Stacked bar graphs color coded by neural G0 phase (G0, G1, G2M/S). In contrast to NECs and progenitors, dCPEC subtypes have predominant G0 signatures (blue) and lower G1 fractions (green) than the neurons. 75-div C1b cells are also G0-predominant, but have a higher G1 fraction than other dCPEC subtypes. Some C2 cells have a G2M/S signature (red) due to shared ciliogenesis and M-phase genes. **(B)** Color coding for subtype clusters used in Fig. 5D. Clusters are assigned to their predominant phase unless a second phase exceeds 30%, in which case a cluster is designated and color coded for “dual” phases. **(C)** 2D PCA scatterplots using neural G0 genes. The second PC distinguishes the neurons (left) while the third PC distinguishes the dCPECs (right). **(D)** UMAP as in Fig. 5, but using Seurat cell cycle genes (mouse), color coded for cell cycle phase. The dCPECs and neurons have predominant G0/G1 signatures (red). G1/G2M C2 cells are also evident (blue). **(E-F)** 3D PCA plot of subtype clusters using Seurat cell cycle genes, color coded for subtype or cell cycle phase as in C or D, respectively. The three branches of NEC progeny are evident. **(G-H)** 3D PCA plot of 31-div cell-type clusters using Seurat cell cycle genes, color coded for cell type (G) or cell cycle phase (H). G0/G1 dCPEC and neuron clusters are distinct. **(I)** The same field as in Fig. 5G, without KI67 and with Hoechst, to enable non-dCPEC estimates outside the dCPEC island.

**FIGURE S8:**
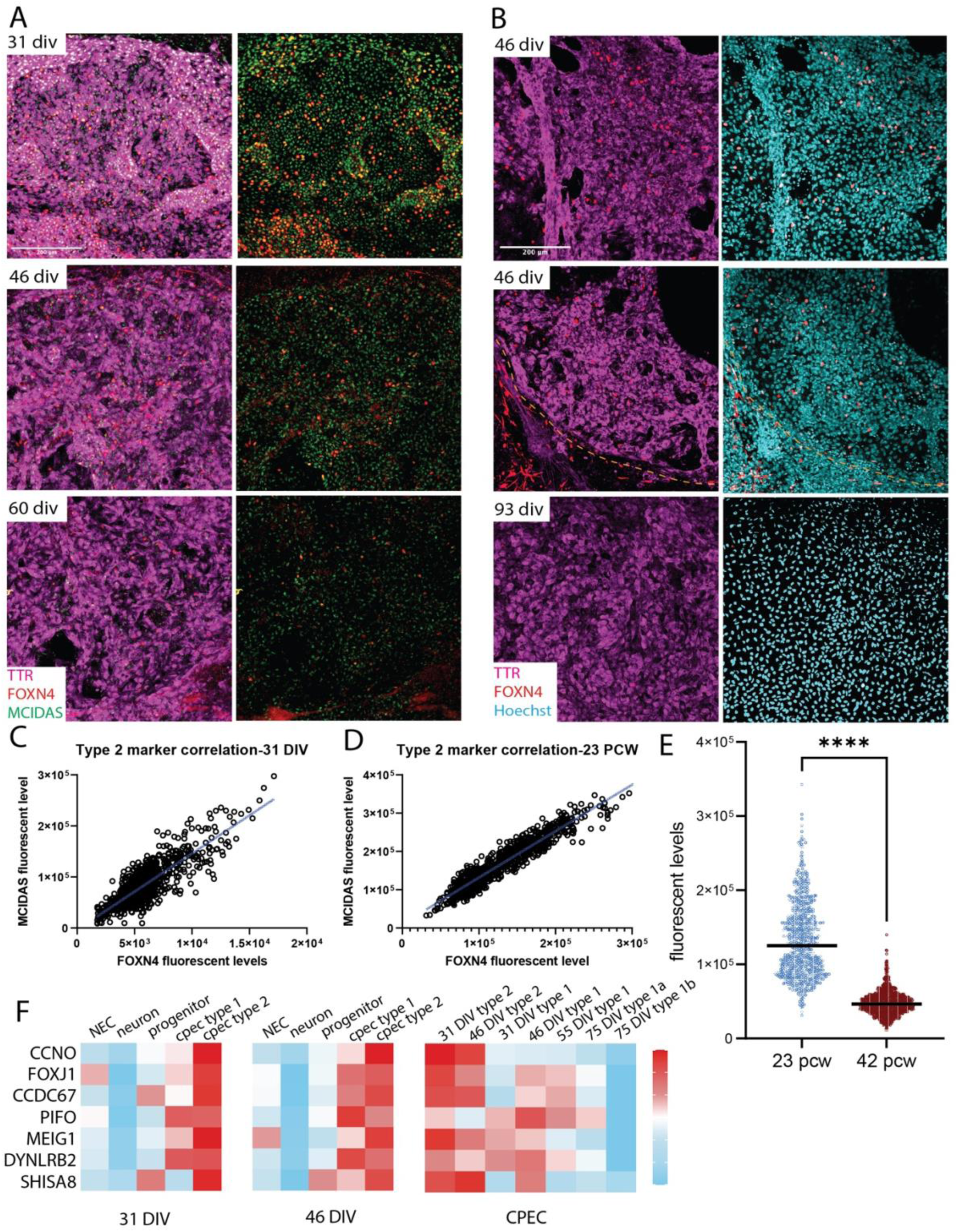
Additional studies on early type 1 and type 2 dCPECs. All images are confocal maximal projections. **(A)** Type 2 dCPECs in parallel cultures (FOXN4-MCIDAS-TTR ICC; 31, 46, and 60 div). FOXN4 (red) and MCIDAS (green) fractional positivities and expression levels decrease over time. **(B)** Lower-power views of the FOXN4 studies shown in Fig. 6F (FOXN4-TTR ICC; 46 and 93 div), dashed lines demarcate dCPEC islands. In addition to the decreasing FOXN4+ C2 fraction, a TTR-negative non-dCPEC population stains for FOXN4 (middle panel). **(C)** Positive linear Pearson correlation of FOXN4 and MCIDAS fluorescence levels in 31-div dCPECs (r=0.8308, p<0.0001; n=1003). **(D)** Positive linear Pearson correlation of FOXN4 and MCIDAS fluorescence levels in 23 pcw ChP tissue (r=0.9486, p<0.0001; n=799). **(E)** Violin plots with medians of FOXN4 fluorescence levels in CPECs *in vivo*, which decrease between 23 and 42 pcw (t-test p<0.0001****; 23-pcw n=799, 42-pcw n=1504). **(F)** Heatmap of C2 markers, including master ciliogenesis genes CCNO and FOXJ1. Compensation for the contraction of C2 or ciliogenesis-enriched cells is not evident.

**FIGURE S9:**
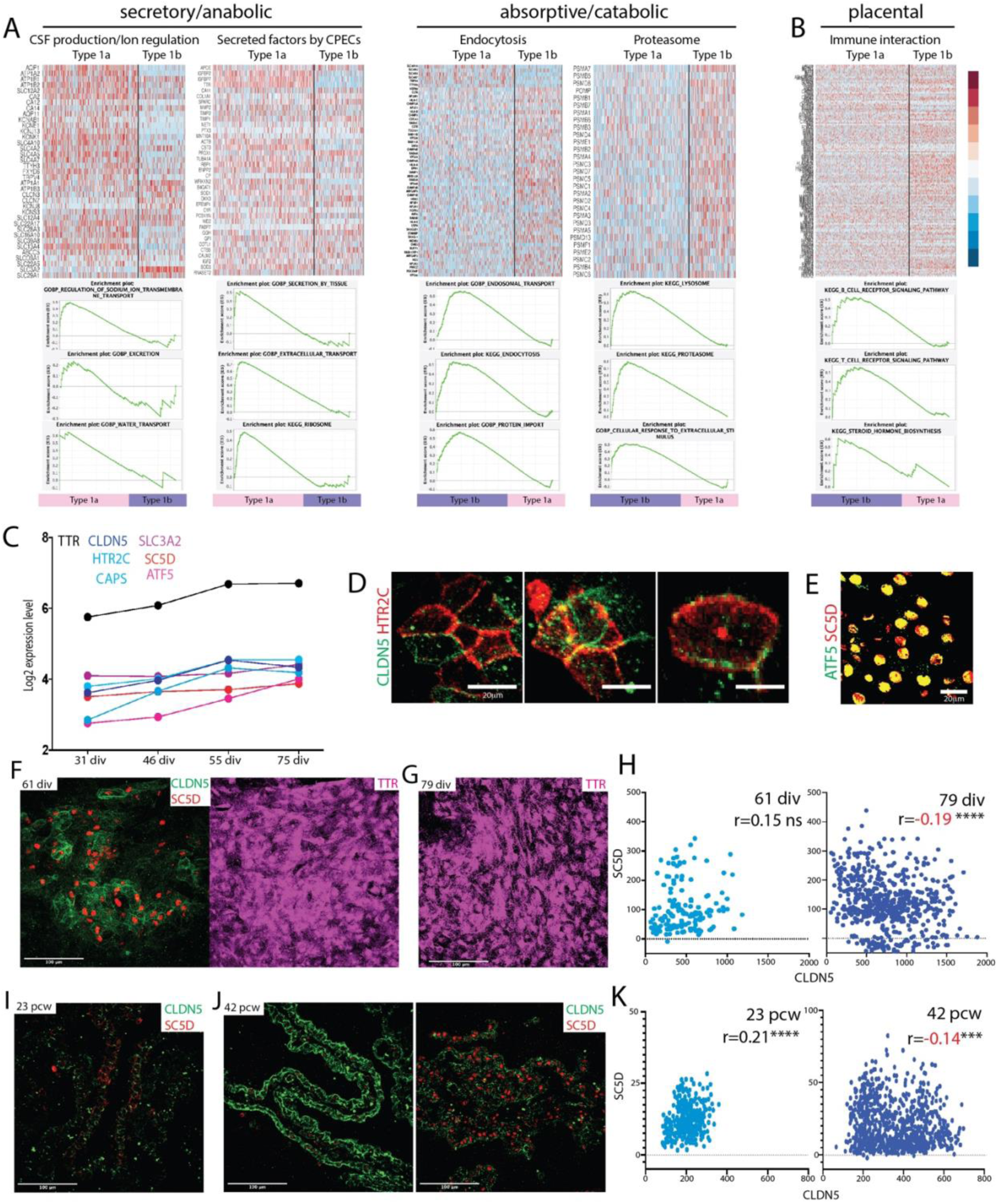
Additional studies on late type 1a and type 1b dCPECs. (A) Secretory/anabolic (left) and absorptive/catabolic pathways (right) enriched in C1a and C1b cells, respectively. (Upper) Heatmaps of manually-curated secretory/anabolic (CSF production/ion regulation, secreted factors) and absorptive/catabolic genes (endocytosis, proteasome). (Lower) GSEA enrichment plots for secretory/anabolic (regulation of sodium ion transmembrane transport, excretion, water transport, secretion, extracellular transport, ribosome) and absorptive/catabolic pathways (endosomal transport, endocytosis, protein import, lysosome, proteasome, cellular response to extracellular stimulus). **(B)** Placental functions^86^ enriched in C1b cells. Heatmaps and GSEA enrichment plots for immune interaction genes, B-cell receptor signaling, T-cell receptor signaling, and steroid hormone biosynthesis. **(C)** Expression levels in individual genes used for pairwise correlations in Fig. 7D (C1a markers: CLDN5, HTR2C, CAPS; C1b markers: SLC3A2, SC5D, ATF5). **(D)** ICC validation of CLDN5 as a C1a marker based on membrane colocalization with HTR2C, another C1a marker (75 div). **(E)** ICC validation of SC5D as a C1b marker based on nuclear colocalization with ATF5, another C1b marker (75 div). **(F-H)** C1a-C1b specification *in vitro* (CLDN5-SC5D ICC maximal projections and scatterplots with Pearson correlations; 61 and 79 div). TTR+ dCPECs at 61 div displaying positively-correlated CLDN5-SC5D expression (F); TTR positivity of the 79 div field shown in Fig. 7F-G (G); scatterplots of CLDN5-SC5D coexpression in individual dCPECs, whose correlation trends positively at 61 div, then becomes significantly negative by 79 div (H) (p=0.15^ns^, n=150; p<0.0001****, n=584). **(I-K)** C1a-C1b specification *in vivo* (CLDN5-SC5D IHC maximal projections and scatterplots with Pearson correlations; 23 and 42 pcw). Relatively low and positively-correlated CLDN5-SC5D expression at 23 pcw (I); relatively large fields of CLDN5^hi^-SC5D^lo^ (C1a) and SC5D^hi^-CLDN5^lo^ (C1b) cells at 42 pcw (J); scatterplots of CLDN5-SC5D coexpression in CPECs *in vivo*, whose correlation is significantly positive at 23 pcw, then becomes significantly negative by 42 pcw (p<0.0001****, n=323; p<0.001***, n=812).

